# The RNA-binding selectivity of the RGG/RG motifs of hnRNP U is abolished by elements within the intrinsically disordered region

**DOI:** 10.1101/2024.05.10.593590

**Authors:** Otto A. Kletzien, Deborah S. Wuttke, Robert T. Batey

**Author notes:** To whom correspondence should be addressed. DSW. RTB.

## Abstract

The abundant nuclear protein hnRNP U interacts with a broad array of RNAs along with DNA and protein to regulate nuclear chromatin architecture. The RNA-binding activity is achieved via a disordered ∼100 residue C-terminal RNA-binding domain (RBD) containing two distinct RGG/RG motifs. Although the RNA-binding capabilities of RGG/RG motifs have been widely reported, less is known about hnRNP U’s RNA-binding selectivity. Furthermore, while it is well established that hnRNP U binds numerous nuclear RNAs, it remains unknown whether it selectively recognizes sequence or structural motifs in target RNAs. To address this question, we performed equilibrium binding assays using fluorescence anisotropy (FA) and electrophoretic mobility shift assays (EMSAs) to quantitatively assess the ability of human hnRNP U RBD to interact with segments of cellular RNAs identified from eCLIP data. These RNAs often, but not exclusively, contain poly-uridine or 5’-AGGGAG sequence motifs. Detailed binding analysis of several target RNAs reveal that the hnRNP U RBD binds RNA in a promiscuous manner with high affinity for a broad range of structured RNAs, but with little preference for any distinct sequence motif. In contrast, the isolated RGG/RG of hnRNP U motif exhibits a strong preference for G-quadruplexes, similar to that observed for other RGG motif bearing peptides. These data reveal that the hnRNP U RBD attenuates the RNA binding selectivity of its core RGG motifs to achieve an extensive RNA interactome. We propose that a critical role of RGG/RG motifs in RNA biology is to alter binding affinity or selectivity of adjacent RNA-binding domains.

## Introduction

Heterogeneous nuclear ribonucleoprotein U (hnRNP U) is a key component of the nuclear scaffold that bridges DNA, RNA and proteins in which its interactions with numerous nuclear RNAs is conferred through a C-terminal domain (U-CTD) comprising two RGG/RG motifs.^1, 2^ RGG/RG motifs are amongst the most common RNA-binding domains (RBDs) in the human proteome, found in >1,700 human proteins by some estimates.^3, 4^ The term RGG/RG motif encompasses a large number of arginine– and glycine-rich regions that contain at least two copies of Arg-Gly-Gly or Arg-Gly motifs, with up to four amino acids bridging each motif.^4, 5^ Due to their low-complexity nature and relative lack of aliphatic amino acids, RGG/RG motifs are frequently located within intrinsically disordered regions.^5, 6^ Although they also interact with other proteins and various macromolecules,^5, 7–11^ RGG/RG motifs are often cited as RNA-binding domains due to numerous studies demonstrating direct RNA recognition by RGG/RG motifs from various proteins, particularly when located adjacent to canonical RNA-binding domains such as the RRM or KH domains.^12–16^

Despite their abundance, a detailed understanding of RNA recognition by RGG/RG motifs remains elusive, likely due to the complexity of studying recognition by intrinsically disordered regions. However, there are several examples of RNA-protein interactions involving RGG/RG motifs that have been structurally characterized. The first is the interaction between the RGG/RG motif of FMRP and SC1, an RNA derived from *in vitro* selection against full-length FMRP protein.^14, 17, 18^ In this complex, the RGG/RG motif occupies a distorted major groove adjacent to a junction between a dsRNA and a G-quadruplex, with arginine residues making base-specific contacts to base pairs within the duplex. This interaction mode has suggested that G-quadruplex motifs are the preferred target of RGG/RG motifs.^19–23^ RGG/RG-mediated interactions of proteins with DNA and RNA G-quadruplexes serve biological roles that include regulation of telomere function,^24, 25^ folding and unfolding of G-quadruplexes^26, 27^ and repression of transcription.^28^ While many RGG/RG proteins bind RNA G-quadruplexes with high affinity,^15,18, 29^ RNAs lacking G-quadruplexes can be bound with comparable affinities,^15^ suggesting a broader RNA-binding mode that is not intrinsically specific for G-quadruplex structure.

Apart from the FMRP-SC1 complex, structural information is available on two additional RGG/RG-RNA complexes: an RGG/RG motif from FUS contacting the minor groove of a model RNA stem-loop,^30^ and an RGGR motif from SF3A1 entering the major groove and forming several base-specific hydrogen bonds with consecutive G-C base pairs in an RNA stem-loop derived from U1 snRNA.^31^ The range of interaction modes displayed in this limited set of available RGG/RG motif structures and the structural variety of their RNA ligands highlights the flexibility of RGG/RG motifs and suggests that they can accommodate a broad range of RNA targets. While RGG/RG motifs interact with diverse RNAs suggesting a promiscuous RNA binding mode, observations of base-specific contacts in RGG/RG–RNA complexes^18, 31^ indicate a potential to recognize specific binding sites within target RNAs.

hnRNP U is an ideal model system to study RGG/RG interactions with RNA as it binds RNA through a C-terminal RGG/RG-rich region and contains no other RBDs.^12, 32^ Furthermore, *in vivo* RNA targets have been identified from eCLIP datasets.^33^ In contrast to previous reports that hnRNP U binds RNA through a single RGG/RG motif,^32^ we found that it uses a large, disordered RBD containing two distinct RGG/RG motifs to achieve full affinity for RNA (**Figure 1**). This full RBD bound a broad range of RNA targets with equilibrium binding affinities <100 nM despite the lack of any shared sequence motif, suggesting a high-affinity, low-selectivity binding mode. Several studies have identified GU-rich sequence motifs near enriched near CLIP-derived RNA-binding sites,^32, 34^ but these motifs have not been shown to be essential for interaction with hnRNP U. More recently, chemical footprinting of hnRNP U bound to a MALT1 pre-mRNA fragment localized binding to a set of GU-rich hairpins.^35^

**Figure 1.**
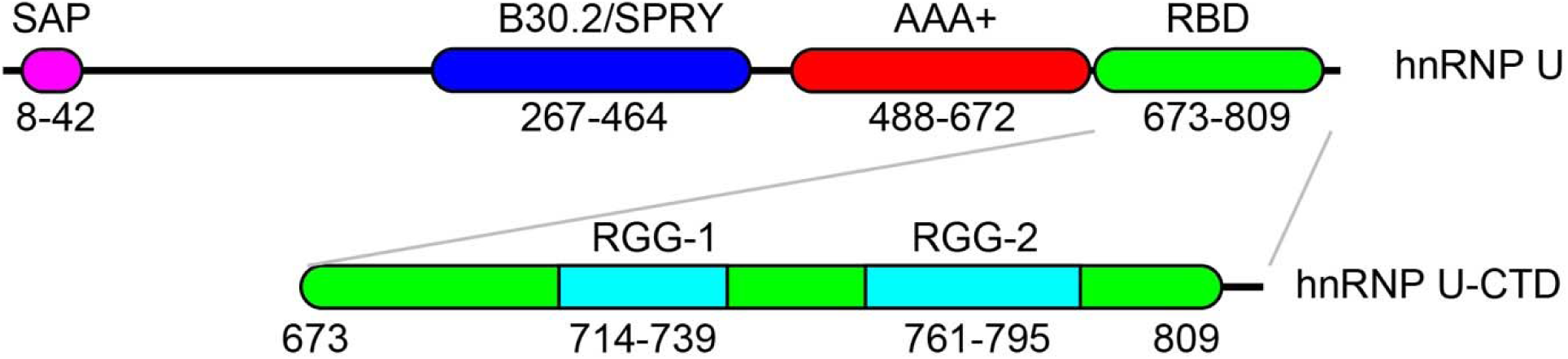
Domain and motif architecture of hnRNP U. The full-length protein, spanning 825 amino acids, is divided into four domains: the DNA-binding SAP domain, the SPRY/B30.2 protein-protein interaction domain, the AAA+ oligomerization domain and the RNA-binding domain (RBD). The RBD domain contains two discrete RGG/RG domains (RGG-1 and RGG-2). The N– and C-terminal boundaries of each domain are denoted below the architecture cartoon.

In this study, we set out to reconcile the varying reports of the RNA-binding selectivity of hnRNP U. The full hnRNP U RBD was determined to have a small preference for G-quadruplex RNAs.^15^ The origins of this preference were probed, and we found that in isolation the first RGG/RG motif and dual RGG/RG motifs show a markedly increased affinity for a model G-quadruplex compared to non-quadruplex RNAs. These different binding behaviors of the isolated RGG/RG motifs and the full RBD strongly suggest that flanking amino acids contribute to RNA recognition by moderating the RGG/RG intrinsic preference for G-quadruplex structures. To assess potential sequence selectivity, the binding of the hnRNP U RBD to a panel of biologically relevant target RNAs identified from eCLIP datasets was compared to the binding to their antisense counterparts. We find that for target RNAs as well as their antisense counterparts, hnRNP U binding affinity increases as RNA length increases up to ∼100 nucleotides (nt), suggesting that the size of its RNA target is more important than its sequence content. Together, the length-dependent binding and lack of observable sequence specificity support that the hnRNP U RBD binds RNA with no requirement for a consensus sequence motif, explaining observations that it interacts with a broad spectrum of nuclear RNAs.^36^

## Results

### G-quadruplex structures are enriched amongst RNA targets of hnRNP U

Because G-quadruplex motifs have emerged as targets of RGG/RG-containing RNA-binding domains in a number of studies,^19–23^ we investigated whether RNA G-quadruplexes were enriched among the set of 323 RNA targets bound by hnRNP U that we previously identified from eCLIP datasets.^33^ Two separate approaches were taken to address this question: computational prediction of G-quadruplex-forming regions based on sequence composition using pqsfinder,^37^ and comparison with a set of experimentally validated *in vivo* G-quadruplexes identified by rG4-Seq.^38^ To ensure that these regions contained the full sequence required to form a potential G-quadruplexes, the original hnRNP U crosslink sites were expanded by 25 nucleotides (nt) in each direction to yield 52-60 nt RNA sequences for input into pqsfinder (sequences identified in analysis of eCLIP data and associated analysis of G-quadruplex formation is provided in Supplemental Data). Of the 323 sites, 100 were identified as predicted G-quadruplexes by pqsfinder and 21 were identified by rG4-Seq, with 16 sites identified by both methods (**Figure 2A**). Both approaches indicate significant enrichment of predicted G-quadruplexes near hnRNP U crosslink sites (**Figure 2B/C**), as determined by permutation tests with random size-matched genomic regions (p-value <0.001 for each method).^39^ Thus, although they do not appear to be strictly required for interaction with hnRNP U, this analysis suggests that G-quadruplexes may represent a preferred subset of hnRNP U RNA targets.

**Figure 2.**
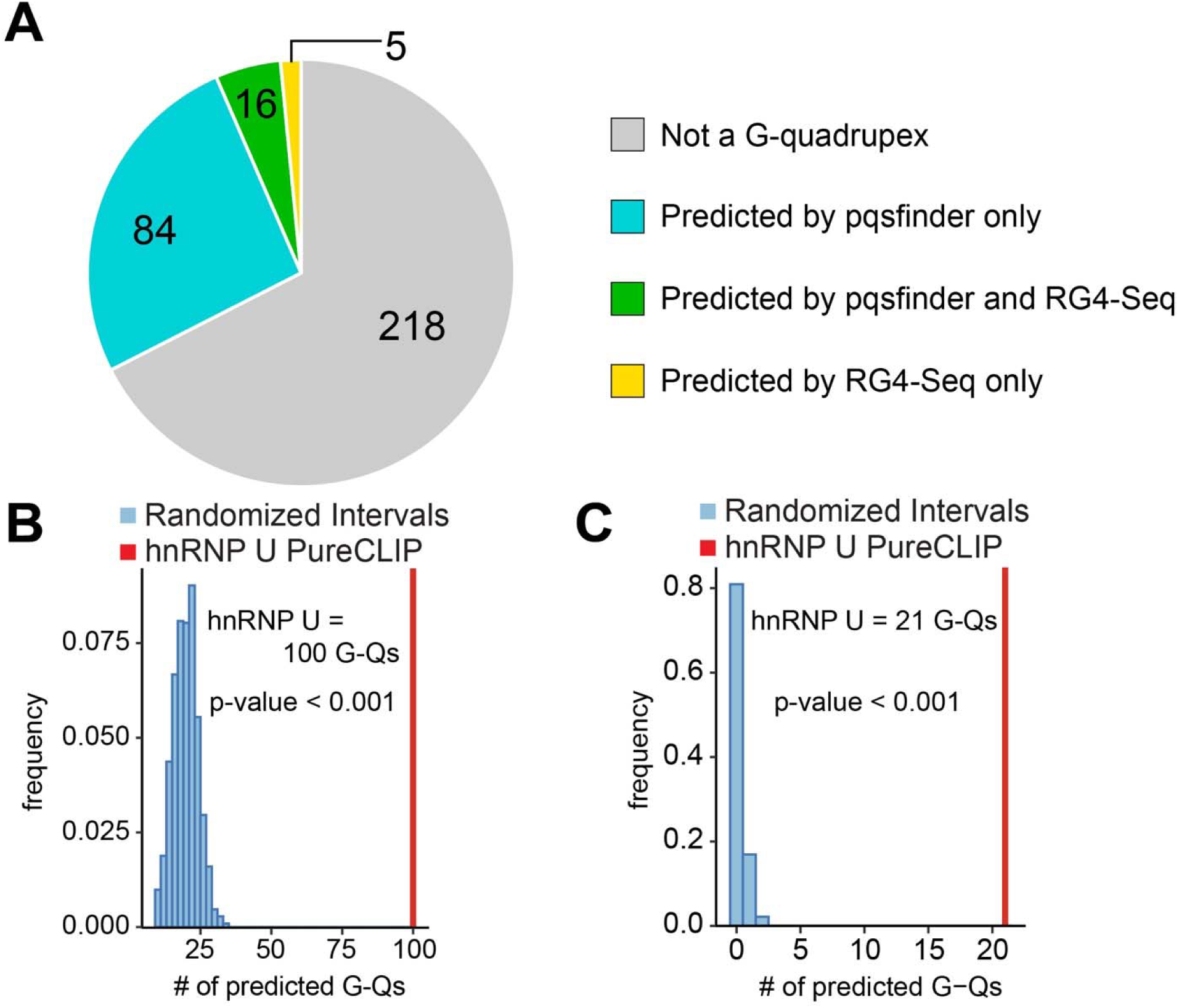
Predicted G-quadruplexes are enriched near hnRNP U PureCLIP sites. (A) Distribution of hnRNP U PureCLIP sites located within 25 nt of predicted G-quadruplexes identified by either pqsfinder ^37^ or RG4-Seq ^38^. (B) Permutation test of predicted G-quadruplexes as assessed by pqsfinder. 1064 sets of size-matched random genomic intervals were compared with hnRNP U crosslink sites. (C) Permutation test of *in vivo* G-quadruplexes as determined by overlap with a previous set of validated G-quadruplexes identified by RG4-Seq. 1064 sets of size-matched random genomic intervals were compared with hnRNP U crosslink sites

Previously, RNA fragments corresponding to eleven hnRNP U crosslink sites were demonstrated to bind the hnRNP C-terminal U domain (Figure 1B; referred to as the “U-CTD” in this work) with sub-micromolar affinities ranging from 60 to 1000 µM.^33^ Seven of these RNA fragments contain regions predicted to form G-quadruplexes by either pqsfinder or rG4-Seq (**Table S1**), suggesting this could be a preferred motif. To experimentally determine whether these minimized RNA fragments form G-quadruplexes under our binding conditions, RNase I nuclease probing was performed under K^+^ and Li^+^ conditions. Because K^+^ strongly supports G-quadruplex formation while Li^+^ does not, G-quadruplexes can be identified by an alteration in secondary structure between the two conditions.^15, 18^ In contrast, A-form helical features in RNA are stabilized by Li^+^ compared to K^+^.^40, 41^ RNase I, which cleaves RNA at the phosphodiester bond 3’-side of single-stranded residues with no sequence preferences, was used to probe the structures of RNAs containing predicted G-quadruplexes.^42^ The seven probed RNAs showed differing propensities to form G-quadruplex structures. The Syncrip RNA fragment displays K^+^-dependent protection of a series of guanine residues, indicating that this RNA contains a G-quadruplex **(Figure 3A-B and Figure S1A)**. Furthermore, this G-quadruplex is located immediately upstream of the hnRNP U crosslink site, suggesting that hnRNP U may directly interact with the G-quadruplex. The same K^+^-dependent change in RNase I reactivity was observed in a smaller 45-nt fragment of the Syncrip RNA (called Syncrip G-quad) that spans the G-rich region, confirming that this small region is sufficient for G-quadruplex formation **(Figure 3C-D and Figure S1B)**. Of the seven survey RNAs fragments, strong RNase I reactivity differences were observed in three RNAs: Syncrip, NORAD, and KCNQ1OT1 and weaker reactivity changes in NPM1 **(Table S1 and Figure S2)**. The G-quadruplexes visualized by RNase I footprinting are located within 5 nt of an hnRNP U crosslink site, further suggesting that they may act as preferred binding sites. The other RNAs (NEAT1, GPI and MALAT1) that were predicted to have G-quadruplex character by pqsfinder and/or RG4-Seq did not exhibit cation-dependent changes in RNase I footprinting, suggesting that these RNAs do not contain this motif.

**Figure 3.**
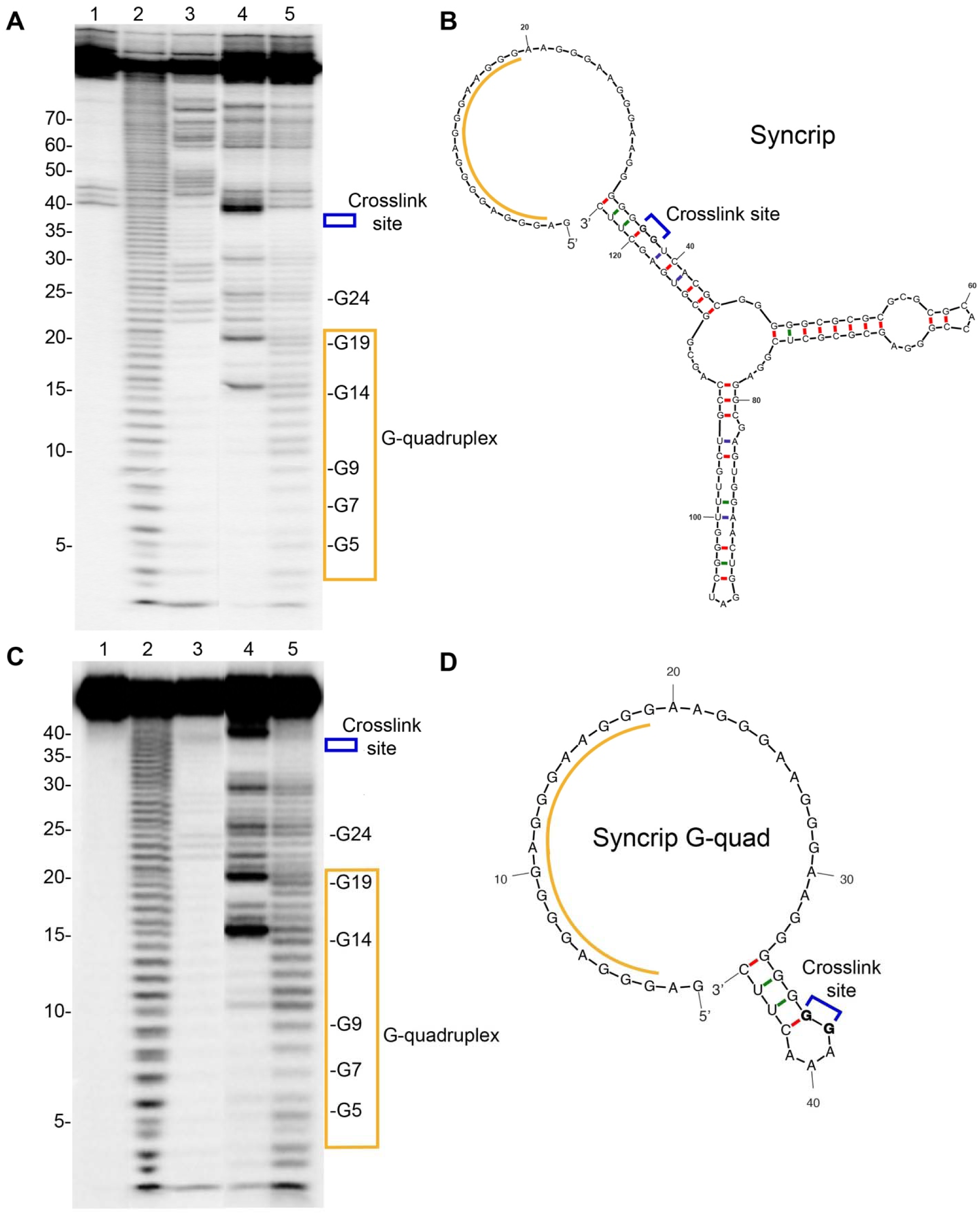
RNA structure probing confirms G-quadruplex formation in Syncrip RNA. (A) RNA structure probing gel of Syncrip RNA. Lanes numbering corresponds to: lane 1, undigested RNA; lane 2, footprinting ladder generated by NaOH cleavage; lane 3, sequencing ladder generated by RNase T_1_ cleavage under denaturing conditions; lane 4; RNase I cleavage in K^+^ buffer; lane 5, RNase I cleavage in Li^+^ buffer. (B) Mfold structure prediction of Syncrip RNA for reference. The region corresponding to the G-quadruplex is labeled in orange and the crosslink site identified by PureCLIP is labeled in blue. (C) RNA structure probing gel of Syncrip G-quad RNA. Lanes are labeled as in part (A). (D) Mfold structure prediction of Syncrip G-quad RNA for reference. The region corresponding to the G-quadruplex is labeled in orange and the crosslink site identified by PureCLIP is labeled in blue. Full gel images for panels A and C shown in Figure S2.

### The hnRNP U RBD shows a modest preference for RNAs containing G-quadruplexes

The Syncrip RNA fragment was selected as a model G-quadruplex-containing RNA to determine whether the U-CTD^33^ interacts specifically with G-quadruplexes. Four Syncrip variants were analyzed: Syncrip RNA **(Figure 2B)** a 45-nt fragment minimized around the G-quadruplex (Syncrip G-quad; **Figure 2D**), and a mutant in which the guanines forming the G-quadruplex were mutated to cytosines (Syncrip GtoC), and a truncated form lacking the region containing G-quadruplex (Syncrip ΔG-quad). U-CTD-RNA binding affinities were measured using a previously established fluorescence anisotropy binding assay.^33^ We found that U-CTD binds both Syncrip RNA and Syncrip G-quad with similar affinities, indicating that the G-quadruplex region encompasses the high-affinity binding site (K_obs_ = 82 ± 15 nM and 80 ± 8 nM, respectively; **Figure 4A**). Deletion of the G-rich region (Syncrip ΔG-quad) or mutation of guanine residues (Syncrip GtoC) are moderately deleterious to binding (350 ± 50 nM and 250 ± 20 nM, respectively), suggesting that hnRNP U shows a modest preference for the G-quadruplex but can interact with RNA sequence outside the G-quadruplex using a weaker binding mode. In addition, the U-CTD binding curves of the RNAs in which the G-quadruplex is abalated are markedly steeper than those for RNAs with a single G-quadruplex, supporting the idea of different binding modes. These steeper transitions may be indicative of multiple proteins binding to non-G-quadruplex RNA, whereas RNAs containing G-quadruplexes exhibit single-site binding behavior.

**Figure 4.**
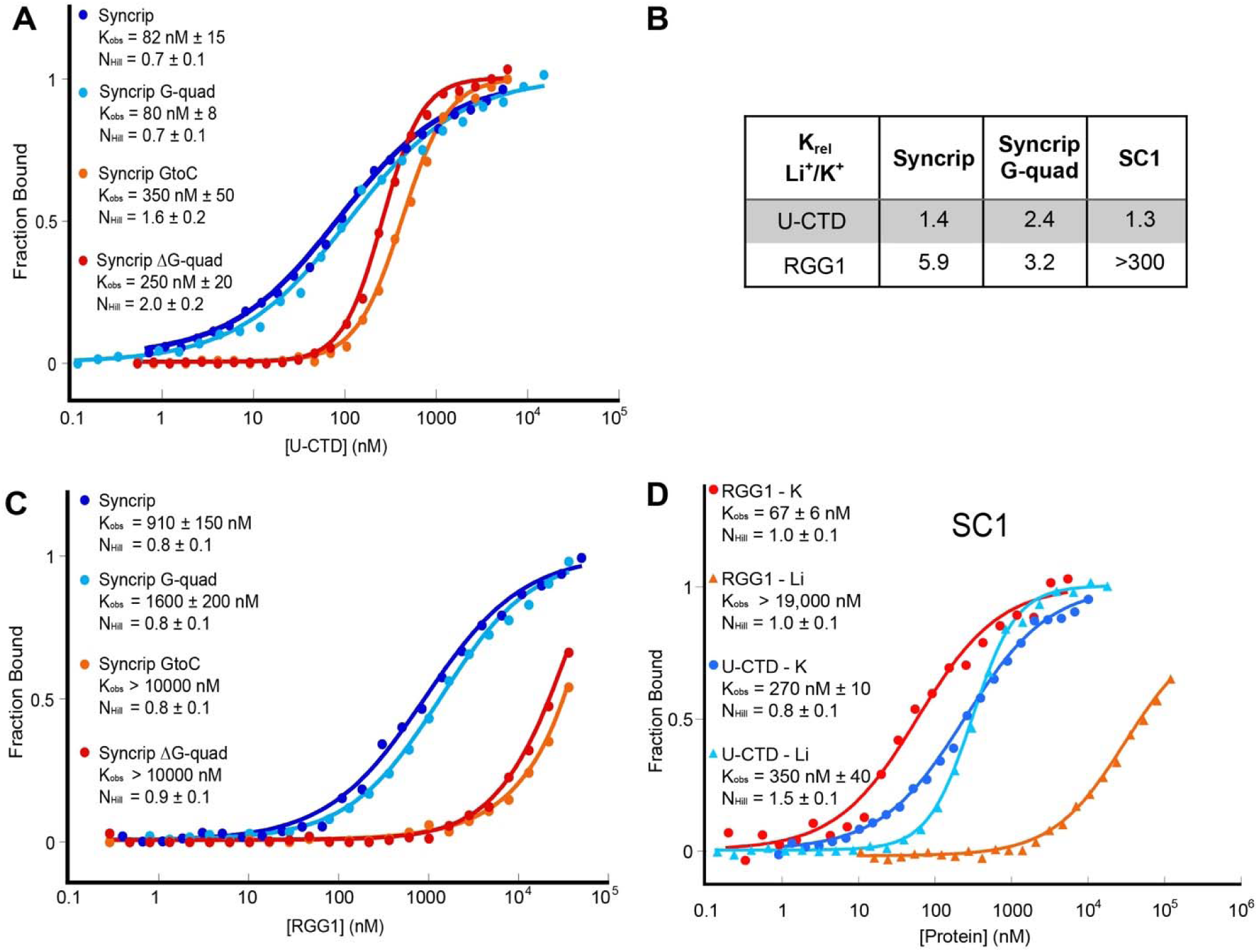
hnRNP U RBD shows a slight binding preference for RNA G-quadruplexes. (A) Representative normalized FA curves of the full hnRNP U RBD binding to the Syncrip variant RNAs. Average K_obs_ values and Hill coefficients are shown with associated standard error of the mean (SEM). (B) Representative normalized FA curves of the minimal RGG/RG motif construct RGG1 bound to the Syncrip variant RNAs. Average K_obs_ values and Hill coefficients are shown with associated SEM. (C) Representative normalized FA curves of RGG1 and U-CTD bound to SC1 in K^+^ and Li^+^ conditions. Average K_obs_ values and Hill coefficients are shown with associated SEM. For RGG1 in K^+^ buffer, the K_obs_ and Hill coefficient of the first transition are shown. (D) Table showing the change in K_obs_ between K^+^ and Li^+^ conditions (K_rel_), for RGG1 and U-CTD bound to Syncrip, Syncrip G-quad, and SC1 RNAs.

To verify the small preference for G-quadruplex structure, the affinities of the hnRNP U-CTD for Syncrip and Syncrip G-quad RNAs were measured in conditions in which K^+^ was replaced with Li^+^. Both RNAs exhibit a very slight decrease in affinity for the hnRNP U-CTD in Li^+^ buffer conditions (K_rel_ values of 1.4 and 2.4 for Syncrip and Syncrip G-quad, respectively, **Figure 4B)**. The loss of affinity induced by unfolding the G-quadruplex was smaller than the loss observed upon deleting or mutating the guanosine-rich region, suggesting that recognition by hnRNP U might be driven by G-rich sequences as well as G-quadruplex structure.

Previous studies using peptides derived from fragile X protein (FMRP) and FUS restricted to the RGG/RG motif sequences have demonstrated a strong preference for G-quadruplex structures.^15, 20, 22^ To assess whether a minimal hnRNP U RGG/RG motif preferentially binds RNAs containing G-quadruplexes, FA assays were performed with a subfragment of the hnRNP U-CTD consisting of the first RGG/RG motif (RGG1, **Figure 1**) and the above set of Syncrip RNA variants. The minimal RGG/RG motif binds both Syncrip and Syncrip G-quad with K_obs_ values of ∼1 µM, approximately 10-fold weaker than the U-CTD **(Figure 4C)**, reflecting the previously observed requirement for the amino acids outside the canonical RGG/RG motif to achieve the highest affinity for RNA.^33^ However, the minimal RGG/RG motif showed significantly weaker affinities for the Syncrip GtoC and Syncrip ΔG-quad RNAs (>10 µM for each). This is a greater loss of binding affinity than observed for the full RBD, suggesting that the RGG/RG motif has enhanced intrinsic selectivity for the RNAs containing the G-quadruplex. Equivalent FA experiments performed with the minimal RGG/RG motif in Li^+^ buffer conditions show a slightly stronger sensitivity to the G-quadruplex structure, as shown by the larger K_rel_ between K^+^ and Li^+^ conditions **(Figure 4C).**

To support these observations, qualitative electrophoretic mobility shift assays (EMSAs) were performed using the four Syncrip variant RNAs and either a truncated form of the full hnRNP U-CTD that facilitates visualization of RNA-protein complexes by EMSA (U-CTD(Δ16))^33^ or the minimal RGG/RG motif (RGG1). These EMSAs confirmed that U-CTD(Δ16) displays moderate selectivity for the Syncrip RNAs containing the G-quadruplex, but maintains affinity for the G-quad deletion and mutant RNAs **(Figure S3A-B)**. The minimal RGG/RG motif exhibits a weak intermediate band at low protein concentrations with G-quadruplex containing RNAs, but a stronger higher-order complex in all RNAs, supporting a G-quadruplex selective binding mode. **(Figure S3C-D)**. Thus, EMSAs qualitatively support the notion that the isolated RGG/RG motif preferentially binds G-quadruplex RNAs as observed by FA.

To further explore recognition of G-quadruplex structures, we performed FA binding analysis using SC1, an *in vitro*-selected aptamer RNA against fragile X protein.^14, 17, 18^ This RNA been shown to bind RGG/RG motifs from multiple proteins with submicromolar equilibrium binding affinities that are dependent upon its G-quadruplex structure.^14, 15^ Comparison of K_obs_ measurements in K^+^ and Li^+^ buffers reveals that the interaction with a pepide bearing the minimal RGG/RG motif from FMRP is >300-fold weaker upon unfolding of SC1 in Li^+^ buffer, while the interaction with the full RBD is largely unchanged (1.3 fold weaker in Li^+^) **(Figure 4B, D)**. These data support previous observations that the RNA-binding behavior of the RBD of hnRNP U and the isolated first RGG/RG motif differ significantly.^33^ Thus, the above results reveal that the hnRNP U RBD has only a modest preference for some G-quadruplex structures in RNAs identified by eCLIP. Since only a subset of these RNAs contain G-quadruplexes, the other RNAs are likely to use other sequence or structural motifs to recruit hnRNP U.

### eCLIP-derived sequence motifs do not drive RNA recognition by hnRNP U

In our recent analysis of hnRNP U eCLIP data, motif analysis by MEME identified two sequence motifs common to sites that showed high-confidence UV crosslinks: 5’-AGGGAG and a U-rich motif.^33^ To further examine whether the 5’-AGGGAG or U-rich motifs represent preferential hnRNP U binding sites, the previously studied NEAT1 and MTRNR2L6 RNAs were selected as model RNAs that respectively contain these motifs.^33^ These RNAs were altered to perturb each motif by either mutating (NEAT1 with a AGGAA to UCCUU mutation and MTRNR2L6 with a UUU to CCC mutation) or deleting the sequence at the crosslink site (Mfold predicted minimal free energy (MFE) structures shown in **Figure S4)**, and the macroscopic binding affinities (K_obs_) of these RNAs for hnRNP U RBD measured by fluorescence anisotropy (FA) (**Figure 5**). Mutation or deletion of the hnRNP U crosslink site in each of these RNAs had almost no effect on K_obs_, suggesting that the U-rich and 5’-AGGGAG motifs are not directly involved in recognition, despite their selective enrichment in eCLIP experiments.

**Figure 5.**
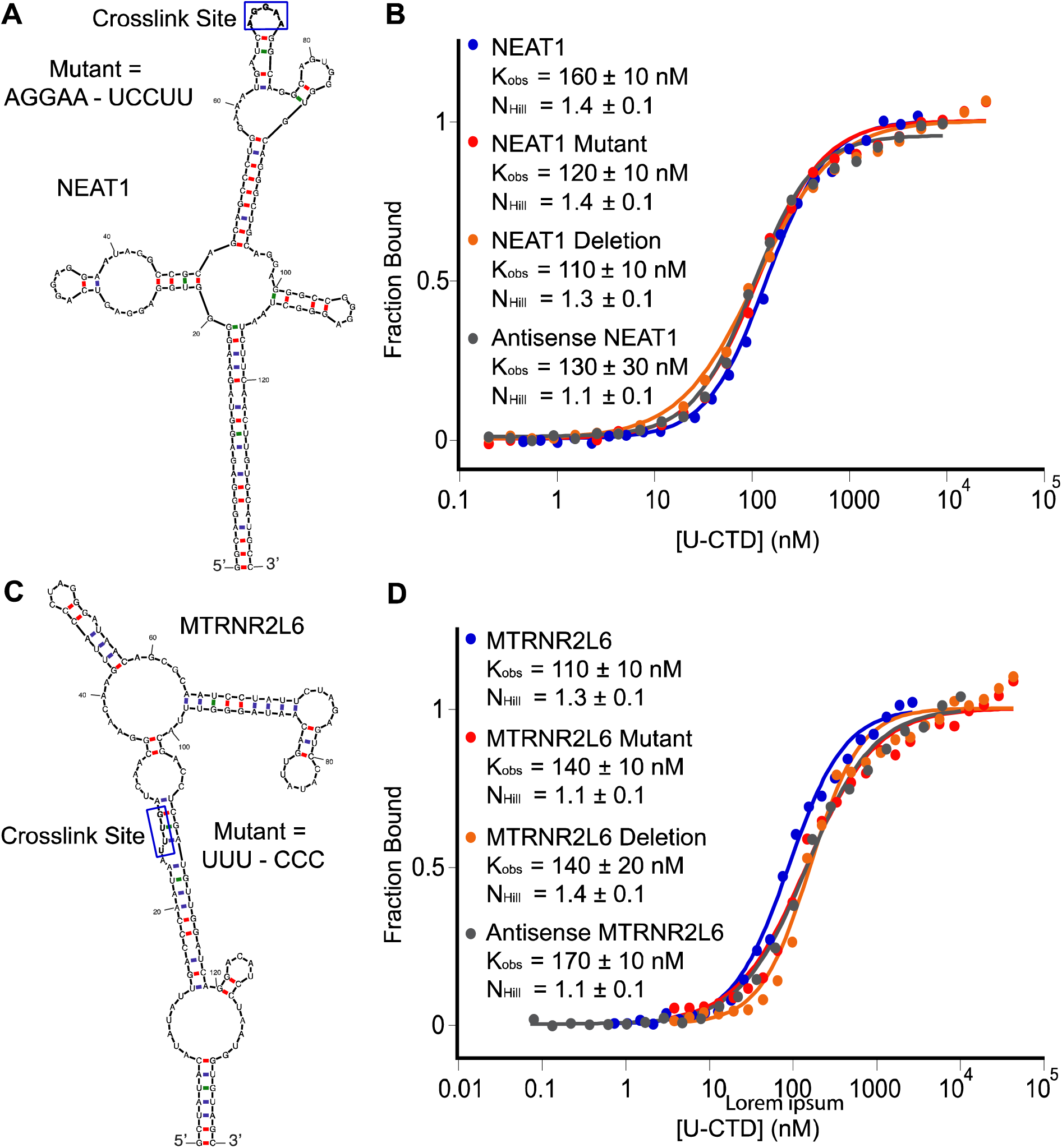
U-rich and AGGGAG motifs are not required for binding. (A) Mfold structure prediction of NEAT1 RNA with the crosslink site highlighted. (B) Representative normalized FA curves of U-CTD bound to NEAT1 variant RNAs. Average K_obs_ values and Hill coefficients are shown with associated SEM. (C) Mfold structure prediction of MTRNR2L6 RNA with the crosslink site highlighted. (D) Representative normalized FA curves of U-CTD bound to MTRNR2L6 variant RNAs. Average K_obs_ values and Hill coefficients are shown with associated SEM.

As a complementary approach to assessing hnRNP U recognition of specific sequence elements within RNAs identified by eCLIP analysis, the antisense complement of seven RNAs resulting from our analysis of eCLIP data were synthesized and tested for binding (**Figure 4B, D and Table 1**). Of the RNAs tested, those containing the 5’-AGGGA and U-rich motifs showed low selectivity for the sense RNA, with the antisense RNAs binding within a two-fold range of K_obs_. Only Syncrip and NPM1, which contain G-quadruplex structures, showed a preference for the sense RNAs over their antisense counterparts (K_rel_ = 4.4– and 3.4-fold, respectively). Together, these findings strongly suggest that hnRNP U does not recognize RNA in a sequence-specific fashion, but rather interacts with a wide range of RNA sequences and structures, consistent with a degenerate specificity model of RNA binding exhibited by a number of proteins.^15, 43, 44^ In this binding mode, also referred to as promiscuous binding, the protein binds a broad range of RNA targets with high affinity and without any apparent sequence preference, although it can exhibit preferences for structural features such as for hairpin loops or dsRNA.^15^

**Table 1:**
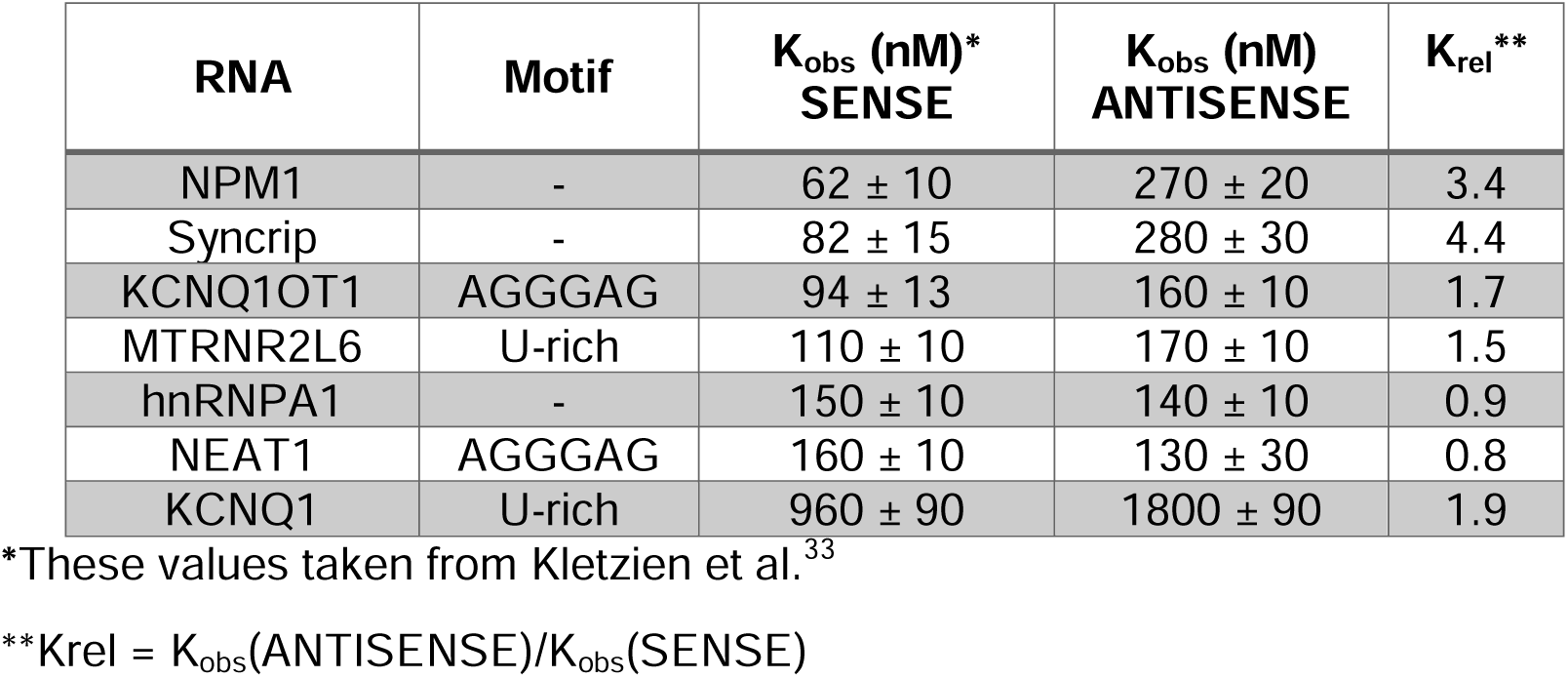
Binding measurements of sense and antisense RNAs to U-CTD.

### U– and G-rich motifs are common eCLIP artifacts

Our data show that hnRNP U binds RNA promiscuously, which raises the question as to why the 5’-AGGGAG and U-rich motifs were highly enriched among its eCLIP crosslink sites. To understand whether these motifs may be crosslinking artifacts, the PureCLIP pipeline was applied to eCLIP datasets from the ENCODE Consortium^45^ for a set of proteins that (*1*) bind RNA solely through one or more RGG/RG motifs, (*2*) proteins that bind RNA through a combination of RGG/RG motifs and other protein domains, and (*3*) proteins that lack RGG/RG motifs and are known to be sequence-specific binders of RNA. MEME^46^ analysis of the regions around the crosslink sites (±10 nt) revealed that U– and G-rich motifs were enriched among proteins regardless of their RBD composition **(Figure 6, Figure S5)**. This finding suggests that U– and G-rich motifs do not reflect specific binding preferences of hnRNP U, in agreement with our data. Furthermore, this analysis suggests these RNA sequence motifs do not reflect general preferences of RGG/RG motifs, as the RNA motifs were identified in proteins that lack RGG/RG motifs altogether. Rather, it is likely that these RNA sequence motifs arise within eCLIP datasets due to UV crosslinking biases. Although UV crosslinking biases towards U-rich sequences have been reported elsewhere^47–49^, the bias towards G-rich sequences that we observed in all sampled eCLIP datasets has not yet been reported to the best of our knowledge.^44^

**Figure 6.**
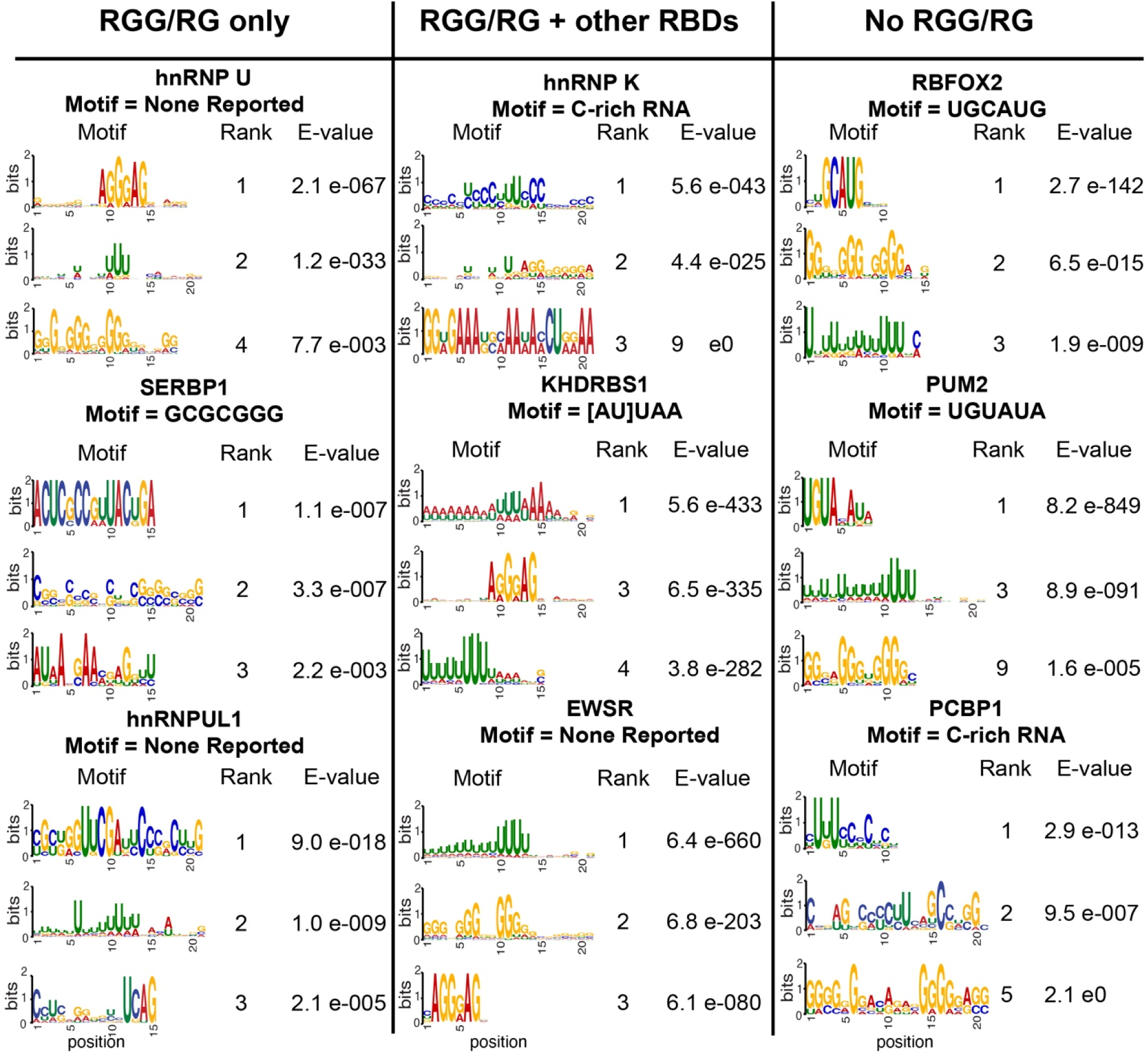
U– and G-rich motifs are frequent eCLIP artifacts. The final PureCLIP sites for each protein were expanded by 10 nt in each direction, then MEME was used to identify enriched motifs. Proteins are separated based on their RBD composition. Previously reported sequence motifs are listed alongside proteins for which motifs have been defined. These motifs are derived from the following sources: SERBP1^59^, hnRNP K^16^, KHDRBS1^60^, RBFOX2^61^, PUM2^62^. PCBP1^63^.

### The hnRNP U-CTD shows length-dependent affinity for RNA

A key characteristic of promiscuous RNA binding is an increase in binding affinity as a function of RNA length due to increased RNA surface area and potential for electrostatically driven interactions, as was previously observed in FUS and PRC2.^43, 50^ To establish whether there is a relationship between the observed affinity and length of the nucleic acid ligand, we used the MTRNR2L6 RNA fragment as a template to generate a systematic set of RNA fragments with lengths varying from 25 to 400 nt (blue series, **Figure 7A**). The K_obs_ of each of these RNA fragments for U-CTD was measured by FA (**Table S2**) and the resulting data plotted as log_10_(K_obs_) versus log_10_(RNA length) (blue series, **Figure 7B**). As RNA length increased from 25 to 100 nt, we observed a nearly linear relationship between K_obs_ and RNA length, which is expected for the RBD binding to a series of sites within each RNA.^51, 52^ As RNA length increases above 150 nt, K_obs_ remains almost constant, indicating that this effect occurs over a limited length regime, and may indicate saturation of the hnRNP U binding interface as was previously observed in PRC2.^43^ This length-dependent change in binding affinity from 25– to 100-nt RNA length mirrored qualitatively by EMSA (**Figure S6**).

**Figure 7.**
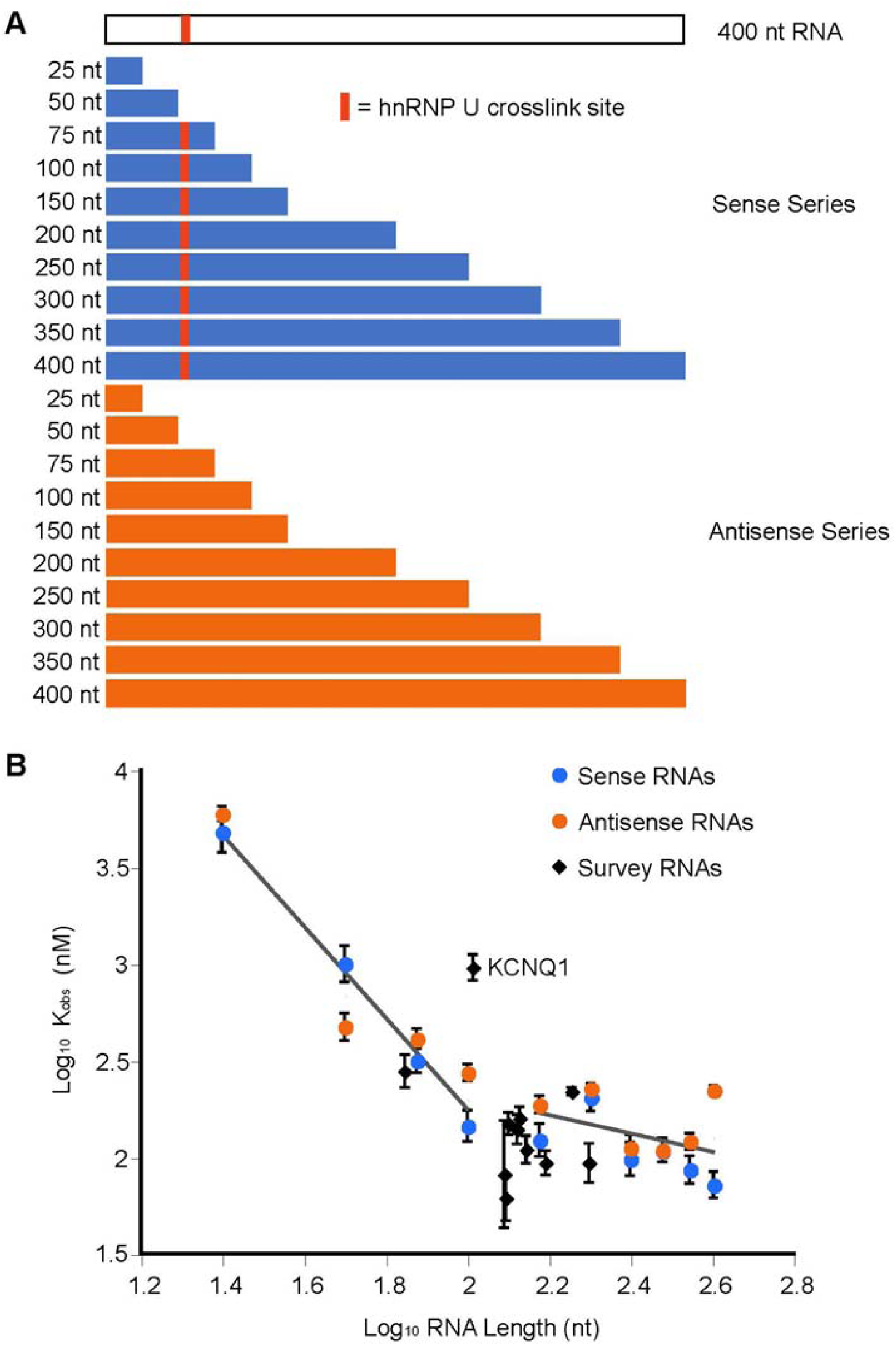
The hnRNP U RBD binds RNA in a length-dependent manner. (A) Schematic of RNA constructs used to test for length-dependent binding. The red line denotes the UUUC crosslink site in the original MTRNR2L6 survey RNA. (B) Plot of log_10_(K_obs_) versus log_10_(RNA Length) for the U-CTD protein fragment binding to the sets of Sense and Antisense MTRNR2L6 RNAs, as well as the set of eleven survey RNAs.^33^ Two lines are shown that fit data between 25 to 100 and 150 to 400 nucleotides in length; these lines intersect at an RNA length of ∼100 nucleotides.

An alternative interpretation to the above observation is that the U-CTD bound all MTRNR2L6 fragment RNAs >150 nucleotides in length with roughly equal affinity because they contain the U-rich crosslink site or a previously unknown sequence feature that drives high-affinity binding. Thus, the observed relationship between affinity and length would reflect two binding modes: a low-selectivity mode for the short RNAs and a site-specific mode for the longer RNAs containing the potential high-affinity binding site. To determine if the presence of such a binding site influences the observed relationship, we synthesized antisense complements for the original set of RNAs, which do not contain the U-rich crosslink site (orange series, **Figure 7A**). Just like the original set of sense-strand RNAs, a plot of log_10_(K_obs_) versus log_10_(RNA length) reveals a strong length dependence for fragments ranging from 25 – 100 nucleotides and independent of length for larger RNAs (orange series, **Figure 7B, Table S2**).

This argues that the breakpoint at ∼100 nucleotides is not due to the addition of a particular binding site, but rather reflects a change once the RNA substrates reach a certain length. In addition, the eleven previously studied hnRNP U target RNAs, with the exception of KCNQ1,^33^ fall within the same K_obs_ range as the MTRNR2L6 fragment RNAs (**Figure 7B**), supporting the idea that hnRNP U binds short RNAs in a length-dependent manner, but longer RNAs within a uniform high-affinity range. Because these target RNAs generally fall within the length-independent regime, their observed binding affinities are not heavily influenced by RNA length. Together, these data further argue that hnRNP U binds RNA in a promiscuous manner wherein binding affinity is determined more by RNA surface area than by sequence content.

### RGG/RG motif selectivity is masked within the context of the full RBD

Since the hnRNP U-CTD binds RNA in a length-dependent manner and shows minimal RNA binding selectivity, we hypothesized that its interactions with RNA may contain a large electrostatic component driven by arginine-phosphate interactions—one of several ways arginine residues interact with RNA.^53^ To test this, binding measurements were performed using the U-CTD against Syncrip G-quad and MTRNR2L6 RNAs over salt concentrations ranging from 25-300 mM KCl. Both RNAs exhibited similar patterns of salt dependence: increasing KCl concentration from 25 to 200 mM did not noticeably affect K_obs_, but the interactions became progressively weaker as KCl concentration was increased from 200-300 mM **(Figure S7A-C)**, similar to the biphasic salt dependence previously observed with FUS.^15^ Since the binding assays in this study were performed in a buffer containing 150 mM KCl, the data reported here is in the binding regime where electrostatic interactions are not dominant. Therefore, although the U-CTD displays promiscuous RNA-binding behavior, this is not reflective of binding dominated by electrostatic interactions but more likely originating from hydrogen bonding to the bases and ribose sugars.^43, 53^

Given the potential importance of electrostatic interactions mediated by the many arginine residues in the RGG/RG motif, the above result is surprising. This may reflect the influence of sequences flanking the RGG/RG motif on determining the nature of the protein-nucleic acid interaction. To probe the extent to which electrostatic interactions drive RNA binding by a single RGG/RG motif, equivalent FA binding assays were performed using the first hnRNP U RGG/RG motif, RGG1, against the Syncrip G-quad and MTRNR2L6 RNAs at salt concentrations ranging from 25-300 mM KCl. In contrast to the hnRNP U RBD, the first RGG/RG motif displays a pronounced salt dependence of K_obs_ **(Figure S7A, D-E)**. Because the binding curves did not saturate at KCl concentrations above 150 mM we could not measure accurate K_obs_ values, but direct comparison of binding curves clearly demonstrates significant loss of affinity with increasing KCl concentration. The interaction with Syncrip G-quad RNA appears to be slightly more salt-tolerant than the interaction with MTRNR2L6 RNA, as assessed by the comparatively smaller shifts from 25 to 150 mM KCl (red & dark green curves, **Figure S6D-E**). These data suggest that compared to the U-CTD, binding of RNA by the RGG1 element in isolation is driven by electrostatic interactions, particularly for non-G-quadruplex RNA (MTRNR2L6). This observation further reinforces our finding that the first RGG/RG motif of hnRNP U does not fully recapitulate the binding behavior of the full hnRNP U RBD and other elements withing the U-CTD are important contributors.

The above data indicate that U-CTD and RGG1 diverge significantly in their RNA recognition features. This suggests that amino acids outside of the first RGG/RG motif modulate/temper the biochemical features of the RGG to make this a better non-selective RNA-binding protein. To further understand how this achieved, a series of U-CTD subfragments was generated, systematically truncating the RNA binding domain towards the first RGG/RG motif **(Figure 8A)**. The binding of these peptides was tested against two RNAs that are discriminated by the RGG but not the full hnRNP U RBD: Syncrip RNA, and its antisense complement that is not predicted to contain a G-quadruplex. Comparison of K_obs_ values for these two RNAs shows the extent to which the presence of the G-quadruplex influences binding by each protein fragment. These data show that as the RBD is minimized to the first RGG/RG motif, selective binding (K_rel_) increases from 3– to 30-fold **(Figure 8B**, **Table 2, Figure S8A-B)**. Notably, the change in binding behavior does not appear to be driven by a specific set of amino acids within the RBD as systematic deletion yields a gradual increase for selective G-quadruplex binding. Thus, the full RBD appears to contribute to high affinity binding of RNAs with or without G-quadruplexes (sense and antisense Syncrip, respectively). However, the core dual RGG/RG motif (hnRNP U(702-793)) maintains strong selectivity for G-quadruplex RNA (K_rel_ = 17), suggesting that sequences flanking the RGG/RG motifs drive a more promiscuous binding mode by the hnRNP U.

**Figure 8.**
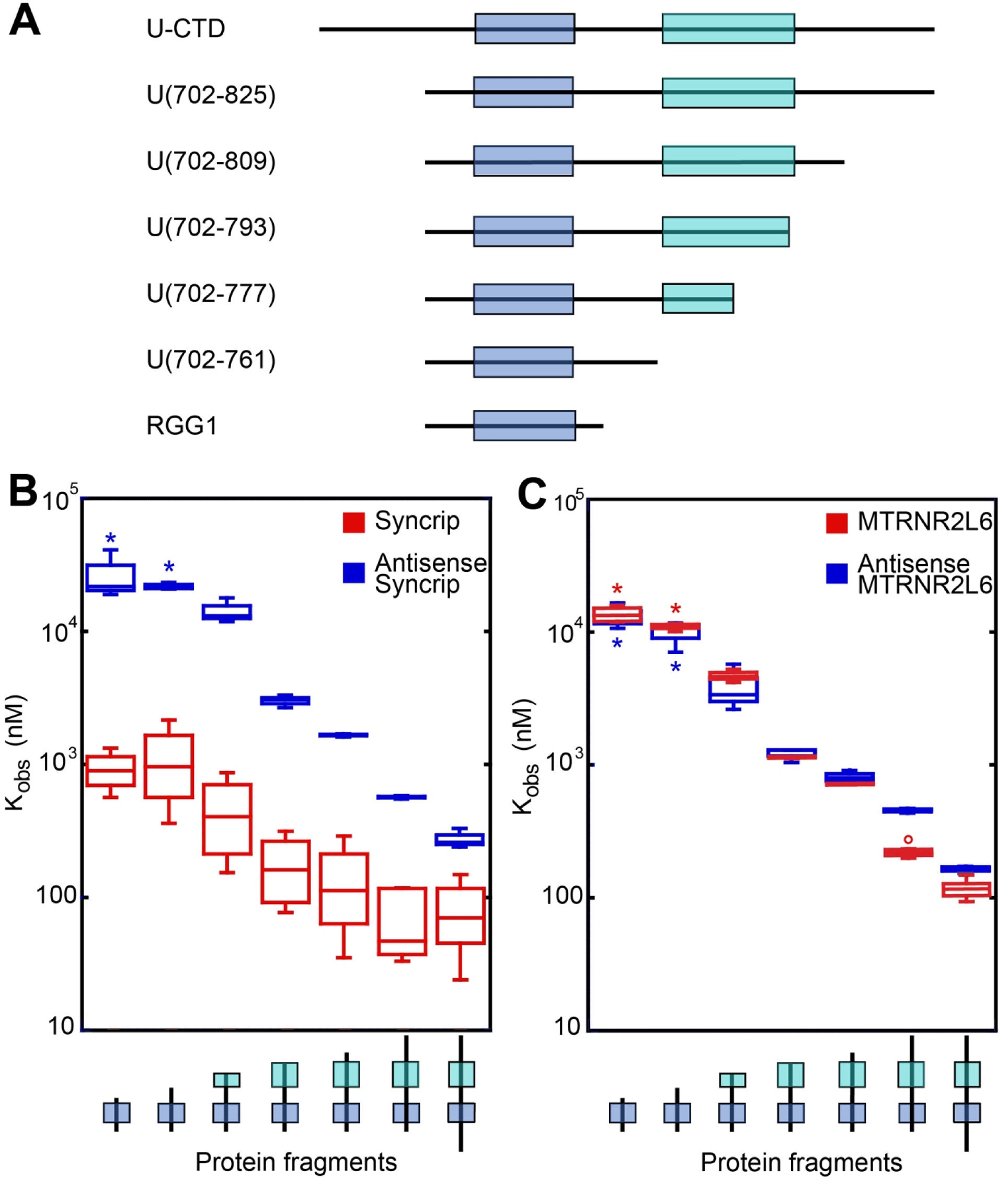
RGG/RG motif selectivity is lost in the context of the full RBD. (A) Schematic of protein fragments. The blue box represents the first RGG/RG motif, and the cyan box represents the second RGG/RG motif. (B) Plot of K_obs_ measurements for each protein fragment vs. Syncrip and Antisense Syncrip RNAs. Asterisks indicate low-end estimates of K_obs_ values from unsaturated binding curves. (C) Plot of K_obs_ measurements for each protein fragment vs. MTRNR2L6 and Antisense MTRNR2L6 RNAs. Asterisks indicate low-end estimates of K_obs_ values from unsaturated binding curves.

**Table 2:** RGG/RG domain selectivity is modulated in the context of the U-CTD.

| Protein | Syncrip<br>$K_{obs}$ (nM) | Anti-Syncrip<br>$K_{obs}$ (nM) | Syncrip<br>$K_{rel}$ | MTRNR2L6<br>$K_{obs}$ (nM) | Anti-MTRNR2L6<br>$K_{obs}$ (nM) | MTRNR2L6<br>$K_{rel}$ |
| --- | --- | --- | --- | --- | --- | --- |
| U(673-825) | 82 ± 15 | 280 ± 30 | 3.4 | 110 ± 10 | 160 ± 10 | 1.5 |
| U(702-825) | 70 ± 19 | 570 ± 10 | 8.1 | 220 ± 10 | 450 ± 10 | 2.0 |
| U(702-809) | 140 ± 50 | 1600 ± 100 | 11 | 710 ± 10 | 810 ± 60 | 1.1 |
| U(702-793) | 180 ± 50 | 3000 ± 200 | 17 | 1100 ± 100 | 1200 ± 100 | 1.1 |
| U(702-777) | 450 ± 160 | 14000 ± 2000 | 31 | 4200 ± 500 | 3900 ± 900 | 0.9 |
| U(702-761) | 1100 ± 400 | > 21000* | >20 | > 11000 | > 9000 | N/A |
| U(702-745) | 910 ± 150 | > 26000 | >20 | > 13000 | > 13000 | N/A |

\* “>” denotes estimated $K_{obs}$ values from unsaturated binding curves.

The same panel of hnRNP U fragments tested for binding against the MTRNR2L6 and antisense MTRNR2L6 RNAs show a decrease in K_obs_ as the RBD is minimized to the first RGG/RG motif **(Figure 8C**, **Table 2, Figure S8C)**. There is no selectivity for the original MTRNR2L6 RNA over its antisense counterpart, consistent with the lack of a G-quadruplex in this RNA. Comparison of these observations strongly suggests that the dual RGG/RG motif in isolation selectively recognizes RNAs containing G-quadruplexes, while the rest of the RBD negates that preference while providing additional binding affinity.

## Discussion

RGG/RG motifs are one of the most abundant RBDs in the human proteome,^3^ but unlike other common RBDs, little is known about their capacity to selectively bind cognate RNA sites. Being intrinsically disordered, RGG/RG motifs can likely adopt numerous conformations to recognize RNA, but it remains unclear whether they can achieve selectivity for RNA targets by recognizing particular sequence/structural features. Prior studies have principally focused on RNA G-quadruplexes as preferred ligands of RGG/RG motifs,^19–23^ but numerous RNAs lacking G-quadruplexes are bound by RGG/RG motifs with biologically relevant binding affinities, suggesting that G-quadruplexes may only represent a subset of RGG/RG targets.^15^ In this work we investigated in detail a set of biologically relevant RNA ligands to understand the RNA-binding selectivity of hnRNP U. Based upon work defining the RNA binding properties of hnRNP U, this study was conducted with a complete domain of the protein that contains all of the sequence requirements for high affinity RNA binding.^33^ Together, these data revealed key features of RNA binding by hnRNP U and have implications for other proteins that use RGG/RG motifs to interact directly with RNA.

### hnRNP U binds RNA with degenerate specificity

The full RNA-binding domain of hnRNP U showed limited selectivity for its RNA targets with only a modest preference for RNAs containing G-quadruplexes. While the hnRNP U RBD binds a select panel of RNAs derived from an analysis of high quality ENCODE eCLIP data,^33^ we also found that the hnRNP U RBD bound antisense RNAs derived from the panel with nearly equivalent affinities, indicating that the protein was likely not preferentially interacting with the identified UV crosslink site. This mode of binding, referred to as degenerate specificity or promiscuous binding, is defined as an RBD having high affinity for numerous RNAs without any apparent consensus sequence motif, but showing a general preference for RNAs with certain secondary structures or combinations of structural elements that are broadly observed in large RNAs. For example, a prior study with the hnRNP U RBD showed a strong preference for RNAs with double-stranded motifs over single-stranded RNA.^15^ Within the panel of eleven RNAs derived from analysis of eCLIP data, ten of these RNAs were bound with macroscopic binding affinities between 60-250 nM, while a single outlier, KCNQ1, was bound with ∼1 μM affinity.^33^ This observation suggests that hnRNP U is capable of discrimination between large RNAs. Interestingly, the KCNQ1 fragment was the only survey RNA with no multi-way junctions in its secondary structure as predicted by Mfold,^33^ which could indicate that complex RNA structures serve as preferred hnRNP U binding sites.

The promiscuous RNA binding observed in this study is consistent with the cellular functions of hnRNP U. Furthermore, a recent study demonstrated that rat hnRNP U directly binds DNA through its C-terminal RNA binding domain, suggesting that its promiscuous binding mode does not strictly discriminate between DNA and RNA ligands.^54^ As a protein that binds DNA, RNA, and numerous other proteins, hnRNP U is proposed to function as a scaffolding protein that regulates numerous cellular processes by bringing various biomolecules into spatial proximity.^55, 56^ Several biological functions of hnRNP U require multivalent interactions, such as attachment of Xist RNA to the inactive X chromosome and maintenance of genome architecture through interactions with hnRNA and the nuclear lamina.^57, 58^ Thus, the degenerate specificity of hnRNP U enables it to bind a broad range of RNAs, giving it the ability to participate in RNA-driven processes without imposing strict sequence requirements and enabling it to serve as a master regulator of RNA biology.

### UV crosslinking bias dominates eCLIP analysis of many RNA binding proteins

The UV-crosslink bias towards uridine and guanosine observed in the analysis of hnRNP U eCLIP datasets is also clear within proteins whose sequence-specific binding sites have been determined independently of eCLIP, including SERBP1,^59^ hnRNP K,^16^ KHDRBS1,^60^ RBFOX2,^61^ PUM2,^62^ and PCBP1.^63^ Although the U– and G-rich biases appear in the sets of motifs identified for these proteins, PureCLIP reliably identifies the established consensus motifs with stronger significance than these background motifs. For sequence specific RNA-binding proteins such as RBFOX2 and PUM2, the specific site is of far greater significance than the U-rich and G-rich background sites (right, **Figure 6**), making identification of the correct site straightforward.

However, crosslink bias can appear within the context of certain canonical motifs, as was observed when the C-rich consensus motifs of hnRNP K and PCBP1. In analysis of eCLIP datasets of these proteins, the C-rich site was observed within the context of the U-rich background motif, despite experimental evidence showing that both proteins binds C-rich sites with high affinity with little selectivity for other nucleotides.^16, 63^ This indicates that for proteins that bind RNA with high sequence specificity, RNAs containing these sites, or closely related sequences, adjacent to U– and G-rich background motifs may improve crosslinking efficiency and thus be over-enriched in the experiment resulting in these nucleotides being included in the binding consensus.

For RBPs with little RNA-binding selectivity, this analysis clearly indicates that the pool of identified motifs will be dominated by crosslinking motifs—a bias that again carries over to the identification of consensus motifs from eCLIP data. A set of proteins without known selectivity yield U-rich and G-rich (or combinations thereof) motifs, as illustrated by EWSR, an RGG/RG motif containing protein that is known to promiscuously associate with RNA (center bottom, **Figure 8**).^64^ Thus, top-ranked U-rich and/or G-rich motifs should be considered a signature of a promiscuous RNA binding protein, which should be should then be validated through biochemical binding analysis. Although eCLIP remains a powerful tool that identifies RNA binding sites for a given RBP, this study further emphasizes that UV crosslinking bias clearly can be a significant impediment to identification of sequence-specific binding preferences.

### The inherent binding properties of minimized RGG/RG motifs are moderated by flanking sequences

In contrast to the full RBD, the isolated dual RGG/RG and first RGG/RG motifs exhibit preferential binding for RNA G-quadruplex structures. While the RBD showed little difference in binding of Syncrip RNA between conditions that promote or disrupt G-quadruplex formation, the first RGG/RG motif showed a 3– to 6-fold higher affinity for Syncrip RNA in the presence of K^+^. Mutation or deletion of the G-quadruplex in Syncrip yield similar losses in affinity for the first RGG/RG motif. Further, a systematic analysis of a set of fragments of RBD that were minimized to the dual and first RGG/RG motifs and tested for binding to a set of four RNAs revealed a marked preference for an RNA G-quadruplex structure within Syncrip RNA.

These results are consistent with other studies of isolated RGG/RG motifs that interact with G-quadruplex structures. A key model system of this interaction has been the FMRP-Sc1 RNA complex.^14^ Sc1 RNA was produced from an *in vitro* selection experiment for RNAs that bound FMRP, which contains a single RGG/RG motif along with several KH domains. Notably, the minimal RGG/RG domain of FMRP binds with the same affinity and selectivity as the full length protein, suggesting that Sc1 captures elements of the FMRP’s preferred cellular RNA targets.^14^ In this study, we observe that Sc1 exhibits 4.5-fold higher affinity for the minimized first RGG/RG motif than the full hnRNP U RBD, in contrast to every other RNA that was assayed. Furthermore, the interaction between Sc1 and the first RGG/RG motif is highly dependent on its G-quadruplex structure; binding to Sc1 was reduced >300-fold in the presence of Li^+^. These data suggest that the Sc1 aptamer may provide a unique binding mode for RGG/RG motifs that does not necessarily recapitulate the properties of native RNA targets of hnRNP U.

Deletion analysis of the RNA-binding domain further suggests that sequences flanking the RGG/RG motif modify its sequence selectivity. In the case of hnRNP U, these flanking sequences abrogate selective binding to G-quadruplex structures to yield a more promiscuous binding mode that favors a spectrum of RNA structures. Thus, these sequences confer a crucial RNA-binding property to hnRNP U. A recent study characterized the RNA-binding properties of PGL-3, a *C. elegans* protein that binds RNA solely through its C-terminal RGG/RG motif and observed widespread RNA binding with low sequence specificity, similar to the promiscuous RNA-binding activity that we report here.^65^ This suggests that the RNA-binding properties of the RBD of hnRNP U are found in other proteins that solely use RGG/RG-containing domains to interact with RNA.

RGG/RG motifs have been described as auxiliary domains as they are often found alongside other canonical RBDs such as RRM, KH, and ZnF domains.^5^ In these cases, the flanking domain dictates the specificity of RNA target binding while the RGG/RG motif confers additional binding affinity. A good example of this is the FUS protein, where the RRM preferentially but weakly binds stem-loop structures, while the adjacent RGG/RG motif significantly enhances protein’s affinity for this site.^30^ This suggests a general model for RGG/RG domains in which the inherent preference of the RGG/RG sequence element for G-quadruplexes is overridden by the specificity of flanking domains that only weakly interact with the target RNA. The function of the RGG/RG domain is not to provide further selectivity for the target, but to increase affinity by interacting with RNA structures adjacent to the target site.

In this fashion, the RGG/RG motif could confer another feature of the protein-RNA interaction—regulation. Modification of the RGG/RG motif or sequestration via interaction with another protein could disrupt or prevent the protein-RNA interaction. Such regulatory functions have been observed in other proteins: recruitment of FUS into stress granules is inhibited by both arginine methylation of its third RGG/RG motif and interactions between the RGG/RG and chaperones including TNPO1 and TNPO3,^66, 67^ while the subcellular localization and stress granule recruitment of another protein, CIRBP, are modulated by arginine methylation and phosphorylation of its RGG/RG motif.^68–70^ Arginines within the first hnRNP U RGG/RG motif are methylated in human cell lines, supporting the idea that this region may be a regulatory center that modulates hnRNP U activity by altering its interactome.^71^ Thus, the RGG/RG motif’s most important contribution to RNA recognition could be its ability to modulate the affinity and specificity of adjacent domains across the multitude of RNA binding proteins in which they are observed.

## Materials and Methods

### Expression and purification of protein constructs

All hnRNP U protein variants were cloned, expressed, and purified as previously described.^33^ Briefly, proteins used in this study were expressed as C-terminal 8xHis-MBP fusions (protein sequences given in Table S3) in *E. coli* Rosetta (DE3) pLysS cells. Following lysis, proteins were purified using a two-column protocol: Ni-NTA (Qiagen) affinity chromatography followed by size exclusion chromatography (HiLoad Superdex 75 pg or 250 pg; GE Life Sciences). Due to the limited solubility of some of the proteins, the 8xHis-MBP domain was retained for all binding experiments to maintain consistency across the study. The 8xHis-MBP tag has no affinity for RNA, as previously established.^15^ For use in fluorescence anisotropy assays performed in Li^+^ conditions, separate batches of protein were purified with solutions in which LiCl was used in place of KCl, at equivalent ionic strengths. After measuring protein concentration by 280 nm absorbance, aliquots were frozen at –80 °C in size exclusion chromatography (SEC) buffer (300 mM KCl or LiCl, 100 mM Tris pH 7.4, 2 mM MgCl_2_, 2 mM DTT, 10% glycerol).

### Preparation, purification, and labeling of RNAs

All RNAs were synthesized by *in vitro* transcription, as previously described (sequences of all RNAs used in this study are given in Table S4).^33^ For use in fluorescence-based experiments, RNAs were labeled at the 3’ end with fluorescein thiosemicarbazide (FTSC), as previously described.^33, 72^ To label RNAs with ^32^P for structure probing assays, unlabeled RNAs were dephosphorylated at the 5’-end by treating with calf intestinal alkaline phosphatase (CIP, New England Biolabs) for one hour at 37 °C. Three phenol/chloroform extractions were performed to remove CIP, then RNAs were ethanol precipitated overnight. Precipitated RNAs were resuspended in Milli-Q water and labeled at the 5’-end with [γ-^32^P]-ATP using T4 polynucleotide kinase (New England Biolabs). Excess [γ-^32^P]-ATP was removed by G-25 MicroSpin column (Cytiva) and eluted into a final volume of ∼50 µL. Labeled RNAs were denatured with 100 µL formamide and 50 µL of formamide dye (95% formamide, 20 mM EDTA, and 0.025% (w/v) bromophenol blue and xylene cyanol), then purified on denaturing acrylamide gels ranging from 6-15% run with 1x TBE buffer (89 mM Tris, 89 mM borate, 2 mM EDTA, pH 8.3), and extracted by crush-and-soak overnight into ∼1 mL of Milli-Q water. RNA extracts were filtered through Corning Co-star 0.22 µm microfuge filters, concentrated to ∼50 µL in Amicon Ultra microfuge concentrator tubes (3-10 kDa MWCO), washed with ∼300 µL Milli-Q water, concentrated to a final volume ∼150 µL, and stored at –20 °C.

### Fluorescence Anisotropy Binding Assays

Fluorescence anisotropy assays were performed as previously described.^33^ In brief, RNAs were refolded by snap-cooling, then diluted to 8 nM concentration in 2x RNA master mix (0.2 mg/mL bovine serum albumin (molecular biology grade, NEB), 0.2 mg/mL yeast tRNA, 10% glycerol) at room temperature. Protein aliquots were thawed and diluted in SEC buffer, centrifuged at high speed for 15 minutes at 4°C, and the supernatant transferred to a new tube. Protein concentrations were determined by 280 nm absorbance, then adjusted to double the starting concentration of the titration. For assays performed in Li^+^ conditions, separate batches of each protein were purified in buffered solutions deplete of K^+^. These aliquots were similarly thawed and diluted in buffered solutions deplete of K^+^. Equal volumes of RNA master mix and protein dilutions were mixed and incubated >45 minutes at room temperature in a 384-well plate before fluorescence measurements were collected. The final reaction conditions were 150 mM KCl (or LiCl), 50 mM Tris pH 7.4, 1 mM MgCl_2_, 1 mM DTT, 10% glycerol, 0.1 mg/mL BSA, 0.1 mg/mL yeast tRNA, 4 nM FTSC-labeled RNA.

Fluorescence anisotropy measurements were taken using a ClarioStar Plus FP plate reader (BMG Labtech). Measured fluorescence intensities were converted to anisotropy using the equation

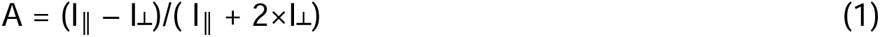

Where A is the fluorescence polarization, I_‖_ is the emission fluorescence intensity parallel to the excitation light and I_┴_ is the emission fluorescence intensity perpendicular to the excitation light. Equilibrium dissociation constants were determined by fitting in Kaleidagraph (Synergy Software) with a one-transition Hill fit. The Hill fit equation used was

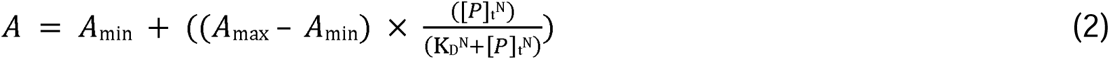

where A is anisotropy, A_min_ is the lower baseline, A_max_ is the upper baseline, [P]_t_ is the total protein concentration, N is the Hill coefficient (referred to as N_H_ elsewhere in text), and K_D_ is the apparent equilibrium dissociation constant.

In cases where binding curves did not appear to saturate at the highest protein concentrations, anisotropy values were normalized to other saturated curves run in parallel using the same protein serial dilution. Each reported K_obs_ is the mean value of at least three separate technical repeats. The standard error of the mean (s.e.m.) was calculated for each K_obs_ and reported.

### Electrophoretic Mobility Shift Assays (EMSAs)

Qualitative EMSAs were performed as previously described.^33^ In brief, protein dilutions and RNA master mix were prepared as described above, except with a higher RNA concentration (24 nM in the master mix for a final concentration of 12 nM). Equal volumes of protein dilution and RNA master mix were mixed and incubated at room temperature for >45 minutes, then 8 µL of each reaction mix was loaded onto a pre-run 6% 29:1 acrylamide:bisacrylamide gel running at 5W in 0.5x Tris-borate running buffer (45 mM Tris-HCl, 45 mM borate, pH 8.2). After running for 35 to 45 minutes, gels were imaged in a Typhoon imager (GE Healthcare) using excitation and emission wavelengths of 495 nm and 520 nm, respectively. The final reaction conditions were 150 mM KCl, 50 mM Tris pH 7.4, 1 mM MgCl_2_, 1 mM DTT, 10% glycerol, 0.1 mg/mL BSA, 0.1 mg/mL yeast tRNA, 12 nM FTSC-labeled RNA.

### Bioinformatic prediction of RNA G-quadruplexes

The set of 323 hnRNP U crosslink sites^33^ were extended by 25 nucleotides in each direction, converted to FASTA files and used as input for pqsfinder with a minimum score cutoff of 25.^37^ All sites with at least one predicted G-quadruplex were considered as “predicted G-quads”. *In vivo* G-quadruplexes were identified from a previous study using RG4-Seq.^38^ Our set of 323 hnRNP U crosslink sites were extended 25 nucleotides in each direction, then bedtools intersect^73^ was used to identify sites conserved between our expanded hnRNP U crosslink sites and the set of G-quadruplexes identified by RG4-Seq (crosslinked in the presence of pyridostatin).^38^ All sites with at least one G-quadruplex were considered “*in vivo* G-quads”.

To assess whether enrichment of G-quadruplexes was statistically significant, permutation tests were performed. A set of 323 genomic regions with 55 nt length was generated by bedtools random to represent random background of similar size, then the process was repeated for a total of 1064 random background distributions. The 1064 sets of random regions were converted to FASTA format and used as input for pqsfinder with the same default settings as the hnRNP U dataset, and the number of predicted G-quadruplexes within each background set was recorded. Student’s t-test was used to assess the statistical significance of G-quadruplex enrichment in the hnRNP U dataset over the 1064 random datasets. The same process was repeated with the “*in vivo* G-quads”, where the 1064 random datasets were searched for previously validated G-quadruplexes using bedtools intersect, then the number of predicted G-quadruplexes within each random dataset was recorded. Student’s t-test was again used to determine statistical significance.

The scripts used to predict G-quadruplex formation and assess statistical significance are available on our GitHub page (https://github.com/bateyLab/RGGbinding_hnRNPu).

### RNA structure probing

To probe the secondary structure of an RNA by RNase I (Thermo Scientific) or RNase T_1_ (Thermo Scientific), 1 µL of labeled RNA was mixed with unlabeled RNA at a final concentration of 1 µM, snap-cooled by heating at 85 °C for 3 minutes and transferring to ice for at least 3 minutes, then mixed with an equal volume of SEC buffer for a final volume of 9 µL and incubated at least 5 minutes at room temperature. 1 µL of 10x RNase I or T_1_, diluted in either 1x KCl or LiCl buffer (150 mM KCl or LiCl, 50 mM Tris pH 7.4, 1 mM MgCl_2_, 1 mM DTT, 5% glycerol), was added to each reaction and allowed to digest at room temperature for 1 minute or 5 minutes, respectively. The 10x stocks of RNase I ranged from 0.16 to 0.00556 U/µL and the 10x stocks of RNase T_1_ ranged from 2 to 0.025 U/µL, depending on the RNA being digested. Alkaline hydrolysis sequence ladders were prepared by adding 1 μL of labeled RNA to 4 µL H_2_O and 10 µL of Alkaline Hydrolysis buffer (50 mM sodium carbonate pH 9.2, 1 mM EDTA) (Ambion). 10 µL of the mix was removed and the remaining 5 µL was incubated at 95 °C for 15 minutes before transferring to ice, adding 3.3 µL of 1M Tris pH 7.4, then adding 25 µL of 8 M urea. RNase T_1_ sequencing ladders were prepared by adding 2 µL of labeled RNA to 5 µL Milli-Q H_2_O, 1 µL of 3 mg/mL yeast tRNA, and 17 µL of 1x RNA Sequencing Buffer (20 mM sodium citrate pH 5, 1 mM EDTA, 7 M urea) (Ambion). The mixture was split across three tubes (9/8/8 µL), which were heated at 85 °C for 3 minutes and cooled at room temperature for 2 minutes. 1 µL of RNase T_1_ at a concentration of 2 U/µL was added to the first tube, then 2 µL of the mixture was transferred to the next two tubes for two successive 1:5 serial dilutions. The second and third dilutions were incubated at 50 °C for 15 minutes before transferring to ice and adding 40 µL of 8 M urea.

Samples were electrophoresed in a 12% 29:1 acrylamide:bisacrylamide gel in 8 M urea and 1xTBE pre-run at 55 W for over 1 hour. 6 µL of each sample was loaded into wells and the gel further electrophoresed for 110-160 minutes. Gels were transferred to Whatman filter paper, dried for 45 minutes in a gel dryer (Bio-Rad), and exposed in a storage phosphor screen (GE Healthcare) for at least 24 hours. Gel cassettes were imaged in a Typhoon imager (GE Healthcare) with the Phosphor setting. Selected gel images were processed using SAFA software to facilitate visualization of predicted G-quadruplex regions.^74^

### PureCLIP analysis of other RNA-binding proteins

Human eCLIP data for several RNA-binding proteins was accessed from the ENCODE Consortium ^45^. Data was accessed for hnRNP K (Experiment ENCSR268ETU, files ENCFF874NKS, ENCFF399CEH, ENCFF405ESF, and Experiment ENCSR828ZID, files ENCFF198ISB, ENCFF019JFZ, ENCFF553XCL), hnRNPUL1 (Experiment ENCSR571VHI, files ENCFF523RZH, ENCFF019BLW, ENCF426AQN, and Experiment ENCSR755TJC, files ENCFF536XVW, ENCFF635FQJ, ENCFF956ELY), SERBP1 (Experiment ENCSR121GQH, files ENCFF943MCZ, ENCFF166ZIY, ENCFF951GRM), KHDRBS1 (Experiment ENCSR628IDK, files ENCFF787ROO, ENCFF931YGS, ENCFF778POI), EWSR (Experiment ENCSR887LPK, files ENCFF842YPC, ENCFF810PWF, ENCFF975NJX), RBFOX2 (Experiment ENCSF756CKJ, files ENCFF537RYR, ENCFF212IIR, ENCFF296GDR, and Experiment ENCSr987FTF, files ENCFF239CML, ENCFF515BTB, ENCFF170YQV), and PUM2 (Experiment ENCSR661ICQ, files ENCFF231WHF, ENCFF786ZZB, ENCFF732EQX). These eCLIP datasets were used as input for the PureCLIP pipeline with the same settings as hnRNP U.^33^ Crosslink sites conserved across biological replicates were identified using bedtools intersect. All eCLIP experiments were performed in HepG2 or K562 cells. Not all of the proteins had available eCLIP data in both cell lines, but in each case, the final set of PureCLIP sites represents sites conserved across all available biological replicates. To identify conserved sequence motifs, each set of PureCLIP sites were extended by four nucleotides in eaxxch direction, converted to FASTA files and used as input for MEME with the following settings: any number of repeats (anr), 1st-order background, motif minimum width = 4 nt, specified strand only.^46^

### CRediT authorship contribution statement

**Otto Kletzien:** Investigation, Formal analysis, Resources, Writing – original draft, Writing – review & editing. **Deborah Wuttke:** Methodology, Resources, Writing – review & editing. **Robert Batey:** Methodology, Resources, Supervision, Writing – review & editing.

## Acknowledgements

We would like to thank Abby Hein for help designing the permutation tests for predicted G-quadruplexes. The authors thank the Shared Instrument Pool (SIP) core facility (RRID: SCR_018986), Department of Biochemistry, University of Colorado Boulder, for the use of the Typhoon FLA 9500 and centrifuges. This work was funded by grants from the National Institutes of Health (R01GM120347 to D.S.W. and R01GM073850 and R35152029 to R.T.B.).

## Appendix A: Supplemenatry data

Supplementary data to this article can be found online at https://xxx.

## Data availability

Data will be made available on request.

