## Supplemental Information for "The RNA-binding selectivity of the RGG/RG motifs of hnRNP U is abolished by elements within the intrinsically disordered region"

**Table of Contents**

Supplementary Tables

Table S1: Survey RNAs containing G-quadruplexes.

Table S2: Apparent binding affinities of length-dependence RNAs for U-CTD.

Table S3: Sequences of proteins used in this study

Table S4: Sequences of RNAs used in this study

Supplementary Figures

Figure S1: RNase I probing of Syncrip and Sycrip G-quad RNAs.

Figure S2: RNase I probing of survey RNAs.

Figure S3: EMSA analysis of U-CTD and RGG1 binding to Syncrip RNA variants.

Figure S4: Mfold structure predictions of NEAT-1 and MTRN2L6 variants.

Figure S5: Top 10 MEME Motifs identified from PureCLIP sites.

Figure S6: EMSAs confirm length-dependent binding by hnRNP U.

Figure S7: U-CTD and RGG1 display varying salt dependencies.

Figure S8: The full hnRNP U-CTD contributes towards binding.

**Table S1:** Features of survey RNAs containing G-quadruplexes using computational and experimental approaches.

| RNA* | Predicted by  pqsfinder | Predicted by  RG4-Seq | RNAse probing |
| --- | --- | --- | --- |
| Syncrip | + | + | + |
| NORAD | + | + | + |
| KCNQ1OT1 | + | - | + |
| NPM1 | + | - | +/- |
| NEAT1 | + | + | - |
| GPI | + | + | - |
| MALAT1 | - | + | - |

Sequences of RNAs given in Table S2.

**Table S2:** Apparent binding affinities of U-CTD for RNAs testing the length dependence of binding.

| RNA Length | SENSE  K_obs_ (nM) | SENSE  N_Hill_ | ANTISENSE  K_obs_ (nM) | ANTISENSE  N_Hill_ |
| --- | --- | --- | --- | --- |
| 25 nt | 4900 ± 700 | 2.2 ± 0.3 | 6000 ± 300 | 3.8 ± 0.2 |
| 50 nt | 1000 ± 100 | 1.5 ± 0.1 | 480 ± 40 | 1.5 ± 0.1 |
| 75 nt | 320 ± 30 | 1.5 ± 0.1 | 410 ± 30 | 1.4 ± 0.1 |
| 100 nt | 150 ± 20 | 0.9 ± 0.1 | 280 ± 20 | 1.7 ± 0.1 |
| 150 nt | 130 ± 10 | 1.0 ± 0.1 | 190 ± 10 | 1.6 ± 0.2 |
| 200 nt | 210 ± 20 | 1.8 ± 0.1 | 230 ± 10 | 1.6 ± 0.2 |
| 250 nt | 100 ± 10 | 1.1 ± 0.1 | 110 ± 10 | 1.1 ± 0.1 |
| 300 nt | 110 ± 10 | 1.6 ± 0.1 | 110 ± 10 | 1.6 ± 0.1 |
| 350 nt | 87 ± 8 | 1.6 ± 0.1 | 120 ± 10 | 1.3 ± 0.1 |
| 400 nt | 73 ± 6 | 1.1 ± 0.1 | 220 ± 10 | 0.7 ± 0.1 |

**Table S3:** Sequences of proteins used in this study.

| **Protein Construct** | **Sequence*** | **Extinction Coefficient**  M^-1^cm^-^ | **pI** |
| --- | --- | --- | --- |
| U-CTD | MHHHHHHHHKIEEGKLVIWINGDKGYNGLAEVGKKFEKDTGIKVTVEHPDKLEEKFPQVAATGDGPDIIFWAHDRFGGYAQSGLLAEITPDKAFQDKLYPFTWDAVRYNGKLIAYPIAVEALSLIYNKDLLPNPPKTWEEIPALDKELKAKGKSALMFNLQEPYFTWPLIAADGGYAFKYENGKYDIKDVGVDNAGAKAGLTFLVDLIKNKHMNADTDYSIAEAAFNKGETAMTINGPWAWSNIDTSKVNYGVTVLPTFKGQPSKPFVGVLSAGINAASPNKELAKEFLENYLLTDEGLEAVNKDKPLGAVALKSYEEELAKDPRIAATMENAQKGEIMPNIPQMSAFWYAVRTAVINAASGRQTVDEALKDAQTNSSSVPGRGSSKKALPPEKKQNTGSKKSNKNKSGKNQFNRGGGHRGRGGFNMRGGNFRGGAPGNRGGYNRRGNMPQRGGGGGGSGGIGYPYPRAPVFPGRGSYSNRGNYNRGGMPNRGNYNQNFRGRGNNRGYKNQSQGYNQWQQGQFWGQKPWSQHYHQGYY | 99,240 | 9.5 |
| U-CTD(∆C16) | MHHHHHHHHKIEEGKLVIWINGDKGYNGLAEKFEKDTGIKVTVEHPDKLEEKFPQVAATGDGPDIIFWAHDRFGGYAQSGLLAEITPDKAFQDKLYPFTWDAVRYNGKLIAYPIAVEALSLIYNKDLLPNPPKTWEEIPALDKELKAKGKSALMFNLQEPYFTWPLIAADGGYAFKYENGKYDIKDVGVDNAGAKAGLTFLVDLIKNKHMNADTDYSIAEAAFNKGETAMTINGPWAWSNIDTSKVNYGVTVLPTFKGQPSKPFVGVLSAGINAASPNKELAKEFLENYLLTDEGLEAVNKDKPLGAVALKSYEEELAKDPRIAATMENAQKGEIMPNIPQMSAFWYAVRTAVINAASGRQTVDEALKDAQTNSSSVPGRGSSKKALPPEKKQNTGSKKSNKNKSGKNQFNRGGGHRGRGGFNMRGGNFRGGAPGNRGGYNRRGNMPQRGGGGGGSGGIGYPYPRAPVFPGRGSYSNRGNYNRGGMPNRGNYNQNFRGRGNNRGYKNQSQGYNQWQQGQ | 83,770 | 9.5 |
| RGG+80 | MHHHHHHHHKIEEGKLVIWINGDKGYNGLAEVGKKFEKDTGIKVTVEHPDKLEEKFPQVAATGDGPDIIFWAHDRFGGYAQSGLLAEITPDKAFQDKLYPFTWDAVRYNGKLIAYPIAVEALSLIYNKDLLPNPPKTWEEIPALDKELKAKGKSALMFNLQEPYFTWPLIAADGGYAFKYENGKYDIKDVGVDNAGAKAGLTFLVDLIKNKHMNADTDYSIAEAAFNKGETAMTINGPWAWSNIDTSKVNYGVTVLPTFKGQPSKPFVGVLSAGINAASPNKELAKEFLENYLLTDEGLEAVNKDKPLGAVALKSYEEELAKDPRIAATMENAQKGEIMPNIPQMSAFWYAVRTAVINAASGRQTVDEALKDAQTNSSSVPGRGSIEGRRGGGHRGRGGFNMRGGNFRGGAPGNRGGYNRRGNMPQRGGGGGGSGGIGYPYPRAPVFPGRGSYSNRGNYNRGGMPNRGNYNQNFRGRGNNRGYKNQSQGYNQWQQGQFWGQKPWSQHYHQGYY | 99,240 | 9.2 |
| RGG+64 | MHHHHHHHHKIEEGKLVIWINGDKGYNGLAEVGKKFEKDTGIKVTVEHPDKLEEKFPQVAATGDGPDIIFWAHDRFGGYAQSGLLAEITPDKAFQDKLYPFTWDAVRYNGKLIAYPIAVEALSLIYNKDLLPNPPKTWEEIPALDKELKAKGKSALMFNLQEPYFTWPLIAADGGYAFKYENGKYDIKDVGVDNAGAKAGLTFLVDLIKNKHMNADTDYSIAEAAFNKGETAMTINGPWAWSNIDTSKVNYGVTVLPTFKGQPSKPFVGVLSAGINAASPNKELAKEFLENYLLTDEGLEAVNKDKPLGAVALKSYEEELAKDPRIAATMENAQKGEIMPNIPQMSAFWYAVRTAVINAASGRQTVDEALKDAQTNSSSVPGRGSFNRGGGHRGRGGFNMRGGNFRGGAPGNRGGYNRRGNMPQRGGGGGGSGGIGYPYPRAPVFPGRGSYSNRGNYNRGGMPNRGNYNQNFRGRGNNRGYKNQSQGYNQWQQGQ | 83,770 | 9.2 |
| RGG+48 | MHHHHHHHHKIEEGKLVIWINGDKGYNGLAEVGKKFEKDTGIKVTVEHPDKLEEKFPQVAATGDGPDIIFWAHDRFGGYAQSGLLAEITPDKAFQDKLYPFTWDAVRYNGKLIAYPIAVEALSLIYNKDLLPNPPKTWEEIPALDKELKAKGKSALMFNLQEPYFTWPLIAADGGYAFKYENGKYDIKDVGVDNAGAKAGLTFLVDLIKNKHMNADTDYSIAEAAFNKGETAMTINGPWAWSNIDTSKVNYGVTVLPTFKGQPSKPFVGVLSAGINAASPNKELAKEFLENYLLTDEGLEAVNKDKPLGAVALKSYEEELAKDPRIAATMENAQKGEIMPNIPQMSAFWYAVRTAVINAASGRQTVDEALKDAQTNSSSVPGRGSFNRGGGHRGRGGFNMRGGNFRGGAPGNRGGYNRRGNMPQRGGGGGGSGGIGYPYPRAPVFPGRGSYSNRGNYNRGGMPNRGNYNQNFRGRGNNR | 75,290 | 9.2 |
| RGG+32 | MHHHHHHHHKIEEGKLVIWINGDKGYNGLAEVGKKFEKDTGIKVTVEHPDKLEEKFPQVAATGDGPDIIFWAHDRFGGYAQSGLLAEITPDKAFQDKLYPFTWDAVRYNGKLIAYPIAVEALSLIYNKDLLPNPPKTWEEIPALDKELKAKGKSALMFNLQEPYFTWPLIAADGGYAFKYENGKYDIKDVGVDNAGAKAGLTFLVDLIKNKHMNADTDYSIAEAAFNKGETAMTINGPWAWSNIDTSKVNYGVTVLPTFKGQPSKPFVGVLSAGINAASPNKELAKEFLENYLLTDEGLEAVNKDKPLGAVALKSYEEELAKDPRIAATMENAQKGEIMPNIPQMSAFWYAVRTAVINAASGRQTVDEALKDAQTNSSSVPGRGSFNRGGGHRGRGGFNMRGGNFRGGAPGNRGGYNRRGNMPQRGGGGGGSGGIGYPYPRAPVFPGRGSYSNRGNYNRGGMP | 73,800 | 8.8 |
| RGG+16 | MHHHHHHHHKIEEGKLVIWINGDKGYNGLAEVGKKFEKDTGIKVTVEHPDKLEEKFPQVAATGDGPDIIFWAHDRFGGYAQSGLLAEITPDKAFQDKLYPFTWDAVRYNGKLIAYPIAVEALSLIYNKDLLPNPPKTWEEIPALDKELKAKGKSALMFNLQEPYFTWPLIAADGGYAFKYENGKYDIKDVGVDNAGAKAGLTFLVDLIKNKHMNADTDYSIAEAAFNKGETAMTINGPWAWSNIDTSKVNYGVTVLPTFKGQPSKPFVGVLSAGINAASPNKELAKEFLENYLLTDEGLEAVNKDKPLGAVALKSYEEELAKDPRIAATMENAQKGEIMPNIPQMSAFWYAVRTAVINAASGRQTVDEALKDAQTNSSSVPGRGSFNRGGGHRGRGGFNMRGGNFRGGAPGNRGGYNRRGNMPQRGGGGGGSGGIGYPYPRAPVFPG | 70,820 | 7.8 |
| RGG1 | MHHHHHHHHKIEEGKLVIWINGDKGYNGLAEVGKKFEKDTGIKVTVEHPDKLEEKFPQVAATGDGPDIIFWAHDRFGGYAQSGLLAEITPDKAFQDKLYPFTWDAVRYNGKLIAYPIAVEALSLIYNKDLLPNPPKTWEEIPALDKELKAKGKSALMFNLQEPYFTWPLIAADGGYAFKYENGKYDIKDVGVDNAGAKAGLTFLVDLIKNKHMNADTDYSIAEAAFNKGETAMTINGPWAWSNIDTSKVNYGVTVLPTFKGQPSKPFVGVLSAGINAASPNKELAKEFLENYLLTDEGLEAVNKDKPLGAVALKSYEEELAKDPRIAATMENAQKGEIMPNIPQMSAFWYAVRTAVINAASGRQTVDEALKDAQTNSSSVPGRGSGGGRGGGHRGRGGFNMRGGNFRGGAPGNRGGYNRRGNMPQRGGGGGG | 67,840 | 7.2 |

*maltose binding protein (MBP) carrier highlighted in green and hnRNP U sequence highlighted in yellow.

**Table S4:** Sequences of RNAs used in this study

| RNA | Length (nt) | Sequence |
| --- | --- | --- |
| Syncrip | 123 | GAGGGAGGGGAGGGAAGGGAAGGGAAGGGAAGGGGGGGUCACGCGGGGGCGCGCGCGCGCACCGGGAGCGCGCUCGGAGGCGAGUGGAACUGGAUCGGGUUUGCUGCCAGCGGCGUGAGCUUC |
| MTRNR2L6 | 139 | GCUAUACAUAUAUUGACCCAAUAAUUUGAUCAACGGAACAAGUUACCCUAGGGAUAACAGCGCAAUCCUAUUCUAGAGUCCAUAUUGACAAUAGGGUUUACGACCUCGAUGUUGGAUCAGGACAUCCUAAUGGUGUAGC |
| NPM1 | 124 | GGGUGGGGAGGCGCGGCGCGGCGUGAGGAGGCCGCAACCGCUGGGAGCACGUGGUUGCCACGUGGUUGGGGGAGGGAGGGGGGUGUGGGCGCUACAUCCGGGACUCACCGGCGUGUUGCCACUC |
| NORAD | 156 | GAGACUAUUCAUUGGGUGUUUGGGGUGGGGGAAGGGGGGGUGGGCAGAGGAGGUAUGCAGGGAGAGGGGUUCUGUGCUCCUGAGAUUAGUUCAGAUGGUCUAACCAUUGUUCUAUAUGUGCAUUUUAGUUAAUAUUGUGUAUUAAAGGAUAAGUCU |
| KCNQ1OT1 | 198 | GGAAGCAACGGCAGCCUGGUCUUUAAGGGUGAGGGAGCACCUUCUCCUGAGAGAGGGAGAGAUAGGUUGGGGGAGGUUGGGUGGGGGUGGGGAGAGCGAGUGAGCAAGCAAGCUAGAAACUUCCUGAAAAGGGAAGUUACUCUAGAAAAGGGUAAGUUCCUUCCAGGCAUCAUGGGCUGGCUUAACCUGGACCAGUUU |
| hnRNPA1 | 126 | GGUUUUAAUGUAGAUUUUUUUUUUUGCACCCCAUGCUGUUGAUUGCUAAAUGUAACAGUCUGAUCGUGACGCUGAAUAAAUGUCUUUUUUUUAAUGUGCUGUGUAAAGUUAGUCUACUCUUAAGCC |
| NEAT1 | 134 | GGCAGGGAGAGGUAGAAGGGGUGGAGGAGUCAGGAGGAAUAGGCCGCAGCAGCCCUGGAAAUGAUCAGGAAGGCAGGCAGUGGGUGCAGGGCUGCAGGAGGGCCGGGAGGGCUAAUCUUCAACUUGUCCAUGCC |
| GPI | 180 | GGGAAUGAGGAGAUGUGUCUUCCAGUGACUUAGGAGGUACGAUCUCUUAUAGGUAUGAGGAUCUGGGUCUGUCAAUGUCUGAGGAGGUAGGAUCUGGCCCUGGUAUGAGAAUCCGGGUCUGUCCAUGUCUGAGCAGGUAGGAUGUGUCCCUGGUAUGAGGAUCCGGGUCUGUCAAUUUCC |
| MALAT1 | 70 | GUUGGCGUGGGGGUGGAGGGGUGAGGUGGGCGCUAAGCCUUUUUUUAAGAUUUUUCAGGUACCCCUCACC |
| KCNQ1 | 103 | GGCCCUCAGUCUACACUCUGUGAUCCACUCAUGAAUAUCUGGUCUGGCUCUGCGUUCCAGUCAGACCCUUUUCUGAGAGGACCACAAGUCAUAGCCCAGGGCC |
| Syncrip G-Quad | 45 | GAGGGAGGGGAGGGAAGGGAAGGGAAGGGAAGGGGGGGAAACUUC |
| Syncrip G-Quad Deletion | 90 | GGGGGUCACGCGGGGGCGCGCGCGCGCACCGGGAGCGCGCUCGGAGGCGAGUGGAACUGGAUCGGGUUUGCUGCCAGCGGCGUGAGCUUC |
| Syncrip G-C mutant | 123 | GAGCCACCCCACCCAACCCAACCCAACCCAACCGGGGGUCACGCGGGGGCGCGCGCGCGCACCGGGAGCGCGCUCGGAGGCGAGUGGAACUGGAUCGGGUUUGCUGCCAGCGGCGUGAGCUUC |
| MTRNR2L6 Crosslink Deletion | 135 | GCUAUACAUAUAUUGACCCAAUAAAUCAACGGAACAAGUUACCCUAGGGAUAACAGCGCAAUCCUAUUCUAGAGUCCAUAUUGACAAUAGGGUUUACGACCUCGAUGUUGGAUCAGGACAUCCUAAUGGUGUAGC |
| MTRNR2L6 Crosslink Mutant | 139 | GCUAUACAUAUAUUGACCCAAUAA**CCCG**AUCAACGGAACAAGUUACCCUAGGGAUAACAGCGCAAUCCUAUUCUAGAGUCCAUAUUGACAAUAGGGUUUACGACCUCGAUGUUGGAUCAGGACAUCCUAAUGGUGUAGC |
| NEAT1 Crosslink Deletion | 129 | GGCAGGGAGAGGUAGAAGGGGUGGAGGAGUCAGGAGGAAUAGGCCGCAGCAGCCCUGGAAAUGAUCGGCAGGCAGUGGGUGCAGGGCUGCAGGAGGGCCGGGAGGGCUAAUCUUCAACUUGUCCAUGCC |
| NEAT1 Crosslink Mutant | 134 | GGCAGGGAGAGGUAGAAGGGGUGGAGGAGUCAGGAGGAAUAGGCCGCAGCAGCCCUGGAAAUGAUC**UCCUU**GGCAGGCAGUGGGUGCAGGGCUGCAGGAGGGCCGGGAGGGCUAAUCUUCAACUUGUCCAUGCC |
| TERRA | 24 | UUAGGGUUAGGGUUAGGGUUAGGG |
| Hairpin No G-Q | 40 | UUAAGAGGCUGAAGCUGAGGCUGAAGCCGAAGCUGAAGCC |
| SC1 | 39 | GGAGCUGCGGUGUGGAAGGAGUGGUCGGGUUGCGCAGCG |
| Antisense Syncrip | 123 | GAAGCUCACGCCGCUGGCAGCAAACCCGAUCCAGUUCCACUCGCCUCCGAGCGCGCUCCCGGUGCGCGCGCGCGCCCCCGCGUGACCCCCCCUUCCCUUCCCUUCCCUUCCCUCCCCUCCCUC |
| Antisense NPM1 | 124 | GAGUGGCAACACGCCGGUGAGUCCCGGAUGUAGCGCCCACACCCCCCUCCCUCCCCCAACCACGUGGCAACCACGUGCUCCCAGCGGUUGCGGCCUCCUCACGCCGCGCCGCGCCUCCCCACCC |
| Antisense NEAT1 | 134 | GGCAUGGACAAGUUGAAGAUUAGCCCUCCCGGCCCUCCUGCAGCCCUGCACCCACUGCCUGCCUUCCUGAUCAUUUCCAGGGCUGCUGCGGCCUAUUCCUCCUGACUCCUCCACCCCUUCUACCUCUCCCUGCC |
| Antisense KCNQ1OT1 | 198 | GGAAAACUGGUCCAGGUUAAGCCAGCCCAUGAUGCCUGGAAGGAACUUACCCUUUUCUAGAGUAACUUCCCUUUUCAGGAAGUUUCUAGCUUGCUUGCUCACUCGCUCUCCCCACCCCCACCCAACCUCCCCCAACCUAUCUCUCCCUCUCUCAGGAGAAGGUGCUCCCUCACCCUUAAAGACCAGGCUGCCGUUGCU |
| Antisense MTRNR2L6 | 139 | GGCUACACCAUUAGGAUGUCCUGAUCCAACAUCGAGGUCGUAAACCCUAUUGUCAAUAUGGACUCUAGAAUAGGAUUGCGCUGUUAUCCCUAGGGUAACUUGUUCCGUUGAUCAAAUUAUUGGGUCAAUAUAUGUAUAG |
| Antisense hnRNPA1 | 126 | GGGCUUAAGAGUAGACUAACUUUACACAGCACAUUAAAAAAAAGACAUUUAUUCAGCGUCACGAUCAGACUGUUACAUUUAGCAAUCAACAGCAUGGGGUGCAAAAAAAAAAAUCUACAUUAAAAC |
| Antisense KCNQ1 | 103 | GGCCCUGGGCUAUGACUUGUGGUCCUCUCAGAAAAGGGUCUGACUGGAACGCAGAGCCAGACCAGAUAUUCAUGAGUGGAUCACAGAGUGUAGACUGAGGGCC |
| 5’SENSE - 25 nt | 25 | GUGCUAAGACUAAACAAGUUUAAGC |
| SENSE - 50 nt | 50 | GUGCUAAGACUAAACAAGUUUAAGCGAAUUACUAUACAUAUAUUGACCCA |
| SENSE - 75 nt | 75 | GUGCUAAGACUAAACAAGUUUAAGCGAAUUACUAUACAUAUAUUGACCCAAUAAUUUGAUCAACGGAACAAGUUA |
| SENSE - 100 nt | 100 | GUGCUAAGACUAAACAAGUUUAAGCGAAUUACUAUACAUAUAUUGACCCAAUAAUUUGAUCAACGGAACAAGUUACCCUAGGGAUAACAGCGCAAUCCUA |
| SENSE - 150 nt | 150 | GUGCUAAGACUAAACAAGUUUAAGCGAAUUACUAUACAUAUAUUGACCCAAUAAUUUGAUCAACGGAACAAGUUACCCUAGGGAUAACAGCGCAAUCCUAUUCUAGAGUCCAUAUUGACAAUAGGGUUUACGACCUCGAUGUUGGAUCAG |
| SENSE - 200 nt | 200 | GUGCUAAGACUAAACAAGUUUAAGCGAAUUACUAUACAUAUAUUGACCCAAUAAUUUGAUCAACGGAACAAGUUACCCUAGGGAUAACAGCGCAAUCCUAUUCUAGAGUCCAUAUUGACAAUAGGGUUUACGACCUCGAUGUUGGAUCAGGACAUCCUAAUGGUGUAGCUGCUAUCAAGGGUUCAUUUGUUCAAUGAUUA |
| SENSE - 250 nt | 250 | GUGCUAAGACUAAACAAGUUUAAGCGAAUUACUAUACAUAUAUUGACCCAAUAAUUUGAUCAACGGAACAAGUUACCCUAGGGAUAACAGCGCAAUCCUAUUCUAGAGUCCAUAUUGACAAUAGGGUUUACGACCUCGAUGUUGGAUCAGGACAUCCUAAUGGUGUAGCUGCUAUCAAGGGUUCAUUUGUUCAAUGAUUAAAGUCCUACGUGGUCUGAGUUCAGACCGGAGCAAGCCAGGUCUGGUACAU |
| SENSE - 300 nt | 300 | GUGCUAAGACUAAACAAGUUUAAGCGAAUUACUAUACAUAUAUUGACCCAAUAAUUUGAUCAACGGAACAAGUUACCCUAGGGAUAACAGCGCAAUCCUAUUCUAGAGUCCAUAUUGACAAUAGGGUUUACGACCUCGAUGUUGGAUCAGGACAUCCUAAUGGUGUAGCUGCUAUCAAGGGUUCAUUUGUUCAAUGAUUAAAGUCCUACGUGGUCUGAGUUCAGACCGGAGCAAGCCAGGUCUGGUACAUGUGUAUUUAAGUCCUCCACACCUCCCAUCUAUAAGUCUUUUUAAAACAAA |
| SENSE - 350 nt | 350 | GUGCUAAGACUAAACAAGUUUAAGCGAAUUACUAUACAUAUAUUGACCCAAUAAUUUGAUCAACGGAACAAGUUACCCUAGGGAUAACAGCGCAAUCCUAUUCUAGAGUCCAUAUUGACAAUAGGGUUUACGACCUCGAUGUUGGAUCAGGACAUCCUAAUGGUGUAGCUGCUAUCAAGGGUUCAUUUGUUCAAUGAUUAAAGUCCUACGUGGUCUGAGUUCAGACCGGAGCAAGCCAGGUCUGGUACAUGUGUAUUUAAGUCCUCCACACCUCCCAUCUAUAAGUCUUUUUAAAACAAAGAAAAUAUGUUUGGUUAAGAGAAGGCUGGAAAUGAGAGAGGGAGUUUUAC |
| SENSE - 400 nt | 400 | GUGCUAAGACUAAACAAGUUUAAGCGAAUUACUAUACAUAUAUUGACCCAAUAAUUUGAUCAACGGAACAAGUUACCCUAGGGAUAACAGCGCAAUCCUAUUCUAGAGUCCAUAUUGACAAUAGGGUUUACGACCUCGAUGUUGGAUCAGGACAUCCUAAUGGUGUAGCUGCUAUCAAGGGUUCAUUUGUUCAAUGAUUAAAGUCCUACGUGGUCUGAGUUCAGACCGGAGCAAGCCAGGUCUGGUACAUGUGUAUUUAAGUCCUCCACACCUCCCAUCUAUAAGUCUUUUUAAAACAAAGAAAAUAUGUUUGGUUAAGAGAAGGCUGGAAAUGAGAGAGGGAGUUUUACUGGACAAAAAAUGUAUAGAGACUUAUGUAACAAUUAUUAUCUGAUAUUAA |
| Antisense - 25 nt | 25 | GCUUAAACUUGUUUAGUCUUAGCAC |
| Antisense - 50 nt | 50 | **G**GGGUCAAUAUAUGUAUAGUAAUUCGCUUAAACUUGUUUAGUCUUAGCAC |
| Antisense - 75 nt | 75 | **G**AACUUGUUCCGUUGAUCAAAUUAUUGGGUCAAUAUAUGUAUAGUAAUUCGCUUAAACUUGUUUAGUCUUAGCAC |
| Antisense - 100 nt | 100 | **G**AGGAUUGCGCUGUUAUCCCUAGGGUAACUUGUUCCGUUGAUCAAAUUAUUGGGUCAAUAUAUGUAUAGUAAUUCGCUUAAACUUGUUUAGUCUUAGCAC |
| Antisense - 150 nt | 150 | **G**UGAUCCAACAUCGAGGUCGUAAACCCUAUUGUCAAUAUGGACUCUAGAAUAGGAUUGCGCUGUUAUCCCUAGGGUAACUUGUUCCGUUGAUCAAAUUAUUGGGUCAAUAUAUGUAUAGUAAUUCGCUUAAACUUGUUUAGUCUUAGCAC |
| Antisense - 200 nt | 200 | **G**AAUCAUUGAACAAAUGAACCCUUGAUAGCAGCUACACCAUUAGGAUGUCCUGAUCCAACAUCGAGGUCGUAAACCCUAUUGUCAAUAUGGACUCUAGAAUAGGAUUGCGCUGUUAUCCCUAGGGUAACUUGUUCCGUUGAUCAAAUUAUUGGGUCAAUAUAUGUAUAGUAAUUCGCUUAAACUUGUUUAGUCUUAGCAC |
| Antisense – 250 nt | 250 | GUGUACCAGACCUGGCUUGCUCCGGUCUGAACUCAGACCACGUAGGACUUUAAUCAUUGAACAAAUGAACCCUUGAUAGCAGCUACACCAUUAGGAUGUCCUGAUCCAACAUCGAGGUCGUAAACCCUAUUGUCAAUAUGGACUCUAGAAUAGGAUUGCGCUGUUAUCCCUAGGGUAACUUGUUCCGUUGAUCAAAUUAUUGGGUCAAUAUAUGUAUAGUAAUUCGCUUAAACUUGUUUAGUCUUAGCAC |
| Antisense – 300 nt | 300 | GUUGUUUUAAAAAGACUUAUAGAUGGGAGGUGUGGAGGACUUAAAUACACAUGUACCAGACCUGGCUUGCUCCGGUCUGAACUCAGACCACGUAGGACUUUAAUCAUUGAACAAAUGAACCCUUGAUAGCAGCUACACCAUUAGGAUGUCCUGAUCCAACAUCGAGGUCGUAAACCCUAUUGUCAAUAUGGACUCUAGAAUAGGAUUGCGCUGUUAUCCCUAGGGUAACUUGUUCCGUUGAUCAAAUUAUUGGGUCAAUAUAUGUAUAGUAAUUCGCUUAAACUUGUUUAGUCUUAGCAC |
| Antisense – 350 nt | 350 | GUAAAACUCCCUCUCUCAUUUCCAGCCUUCUCUUAACCAAACAUAUUUUCUUUGUUUUAAAAAGACUUAUAGAUGGGAGGUGUGGAGGACUUAAAUACACAUGUACCAGACCUGGCUUGCUCCGGUCUGAACUCAGACCACGUAGGACUUUAAUCAUUGAACAAAUGAACCCUUGAUAGCAGCUACACCAUUAGGAUGUCCUGAUCCAACAUCGAGGUCGUAAACCCUAUUGUCAAUAUGGACUCUAGAAUAGGAUUGCGCUGUUAUCCCUAGGGUAACUUGUUCCGUUGAUCAAAUUAUUGGGUCAAUAUAUGUAUAGUAAUUCGCUUAAACUUGUUUAGUCUUAGCAC |
| Antisense – 400 nt | 400 | GUAAUAUCAGAUAAUAAUUGUUACAUAAGUCUCUAUACAUUUUUUGUCCAGUAAAACUCCCUCUCUCAUUUCCAGCCUUCUCUUAACCAAACAUAUUUUCUUUGUUUUAAAAAGACUUAUAGAUGGGAGGUGUGGAGGACUUAAAUACACAUGUACCAGACCUGGCUUGCUCCGGUCUGAACUCAGACCACGUAGGACUUUAAUCAUUGAACAAAUGAACCCUUGAUAGCAGCUACACCAUUAGGAUGUCCUGAUCCAACAUCGAGGUCGUAAACCCUAUUGUCAAUAUGGACUCUAGAAUAGGAUUGCGCUGUUAUCCCUAGGGUAACUUGUUCCGUUGAUCAAAUUAUUGGGUCAAUAUAUGUAUAGUAAUUCGCUUAAACUUGUUUAGUCUUAGCAC |

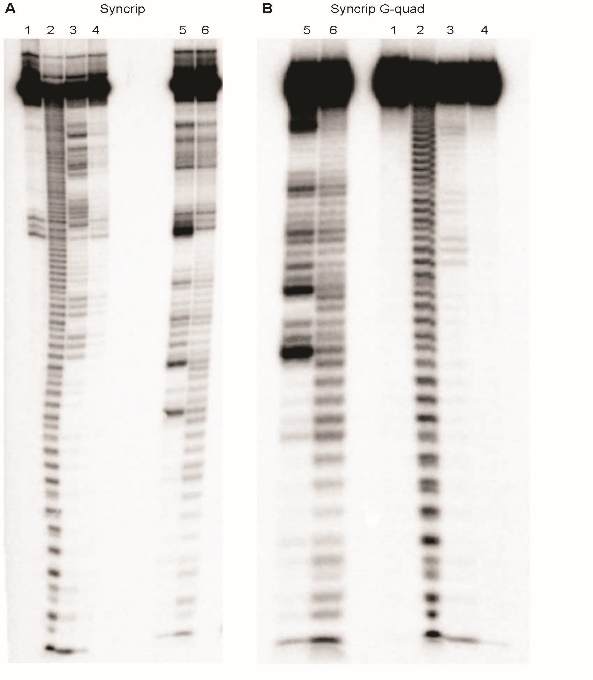

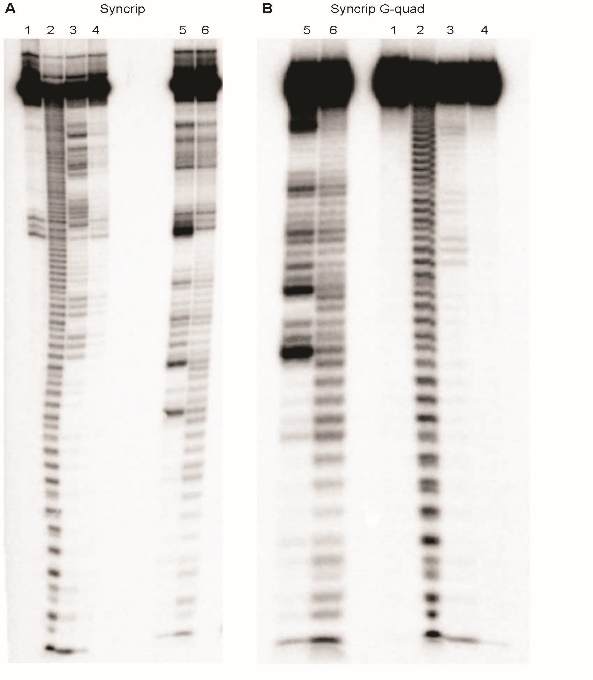
**A	B**

**Figure S1. Raw gels of RNase I probing of Syncrip RNAs.** Probing of (A) Syncrip RNA and (B) Syncrip G-quad. Gel lanes correspond to: lane 1, un-digested RNA; lane 2, nucleotide ladder generated by NaOH cleavage; lanes 3 and 4, sequencing ladder generated by RNase T1 cleavage under denaturing conditions (lane 4 is a 1:5 dilution of RNase T_1_ compared to lane 3); lane 5, RNase I cleavage in K^+^ buffer; lane 6, RNase I cleavage in Li^+^ buffer. Edited gels shown in Figure 3A and 3C.

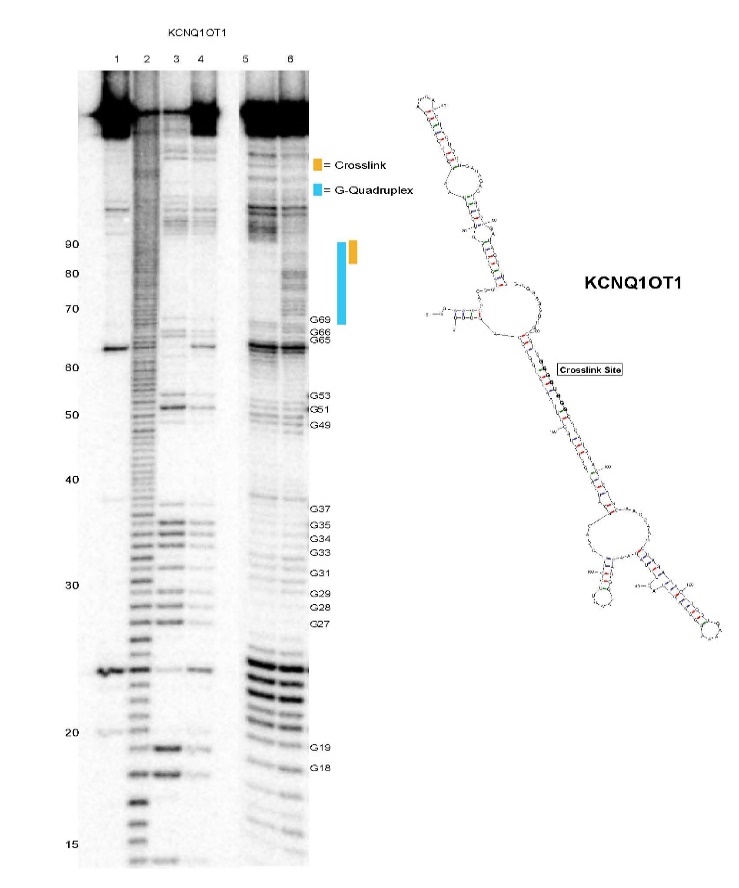

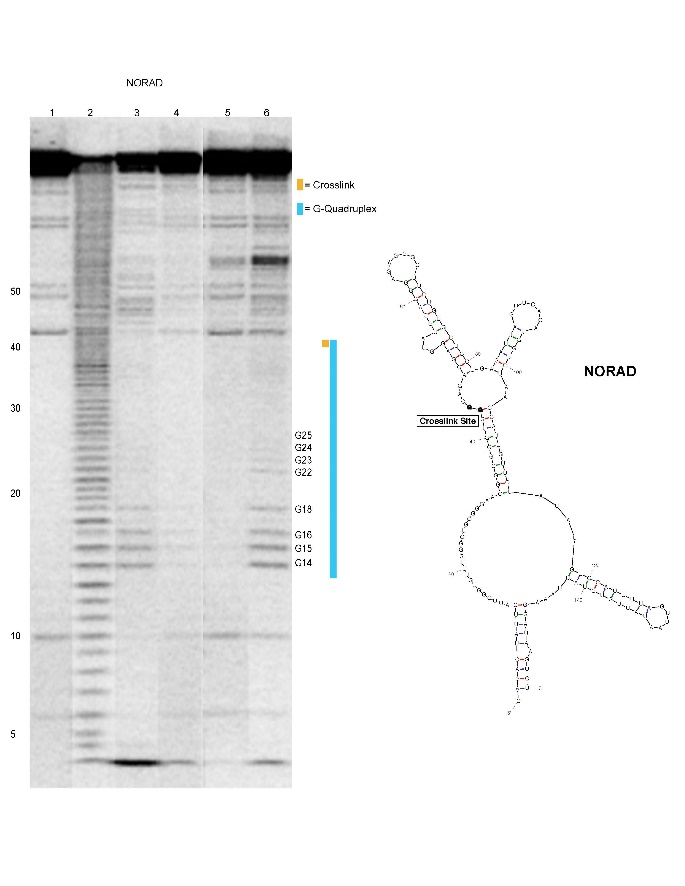
**A	B**

**Figure S2. Raw gels of RNase I probing of survey RNAs.** Continued on next page.

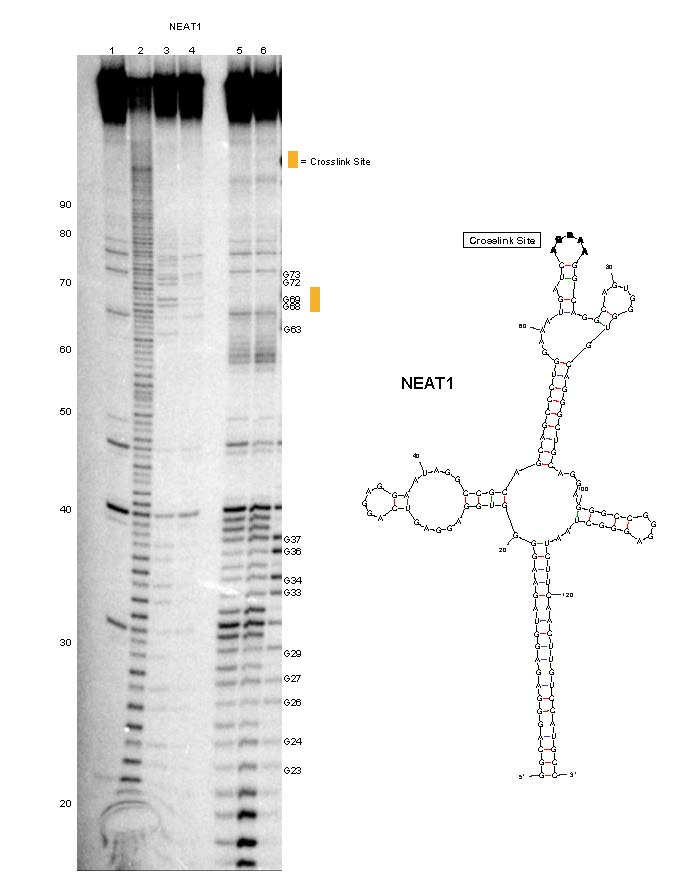
 **C	D**

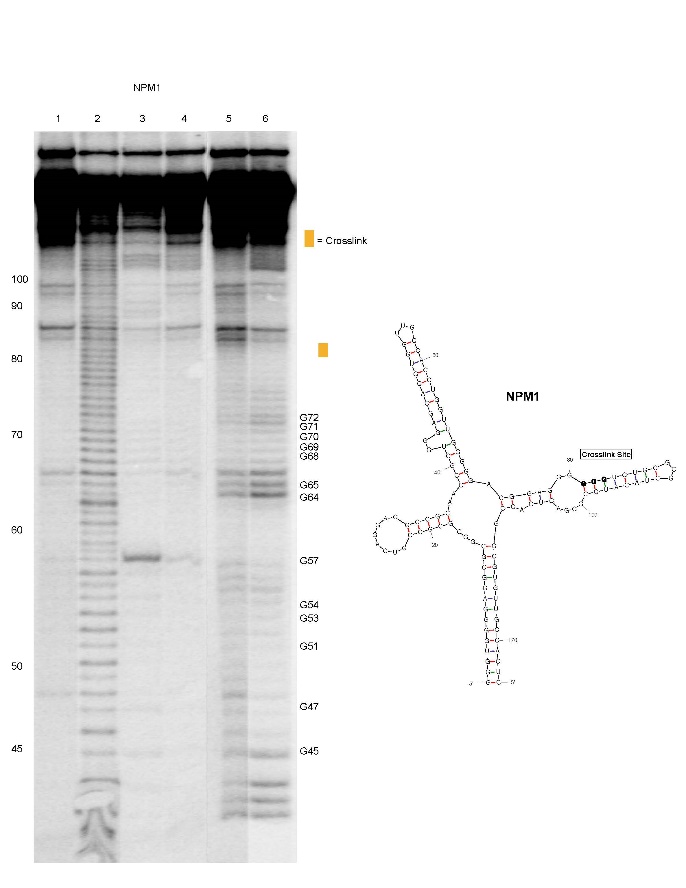

**Figure S2. Raw gels of RNase I probing of survey RNAs.** Continued on next page.

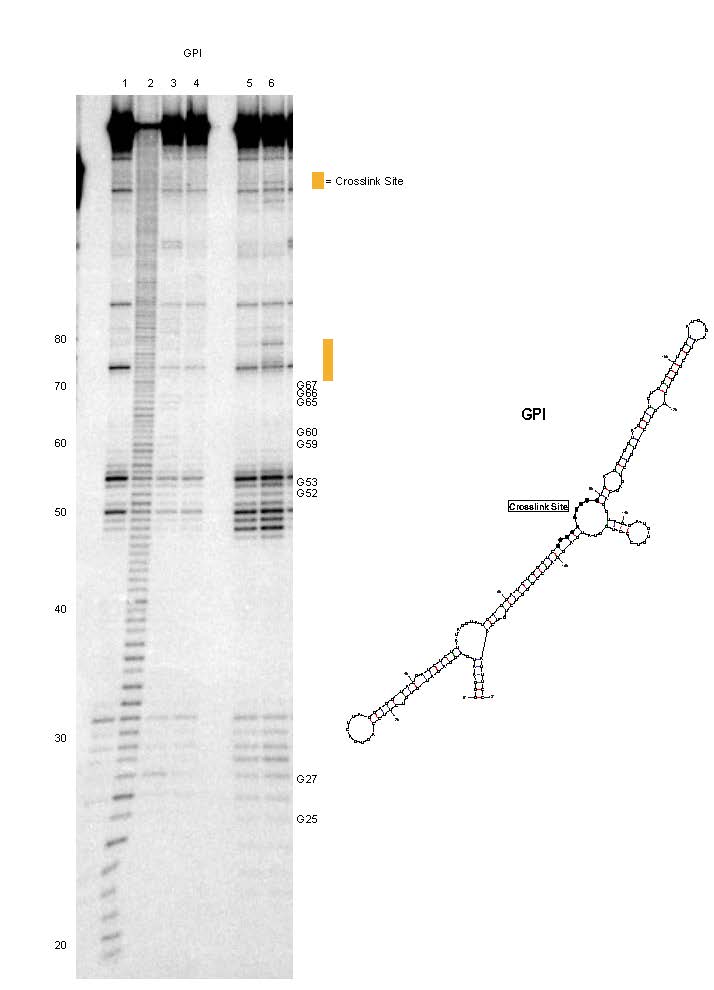
**E	F**

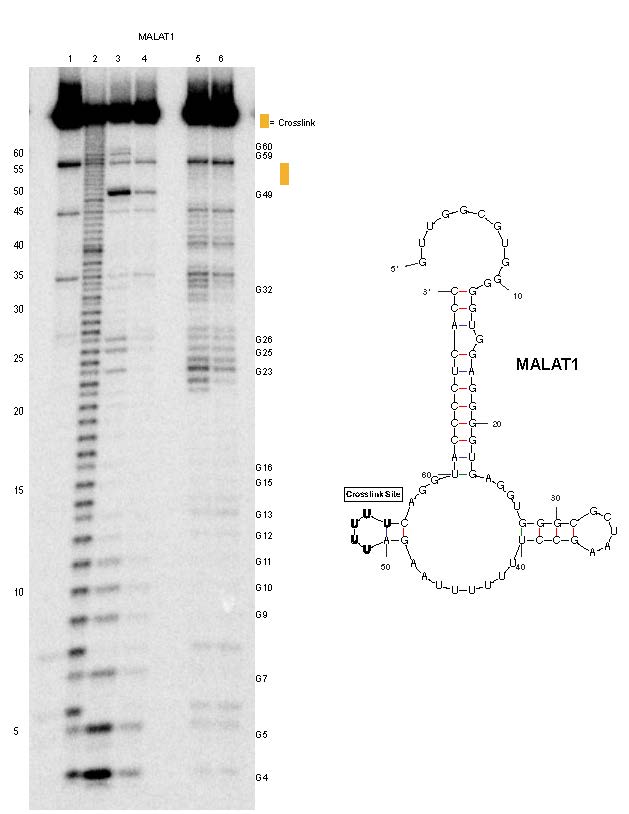

**Figure S2. Raw gels of RNase I probing of survey RNAs.** Probing of (A) NORAD, (B) KCNQ1QT1, (C) NPM1, (D) NEAT1, (E) GPI and (F) MALAT1 RNAs. Orange bars adjacent to gels correspond to the eCLIP crosslink sites and blue bars correspond to G-quadruplex formation. The Mfold-predicted secondary structure of each probed RNA is shown to the right of the gel image. Gel lanes correspond to: lane 1, un-digested RNA; lane 2, nucleotide ladder generated by NaOH cleavage; lanes 3 and 4, sequencing ladder generated by RNase T1 cleavage under denaturing conditions (lane 4 is a 1:5 dilution of RNase T_1_ compared to lane 3); lane 5, RNase I cleavage in K^+^ buffer; lane 6, RNase I cleavage in Li^+^ buffer.

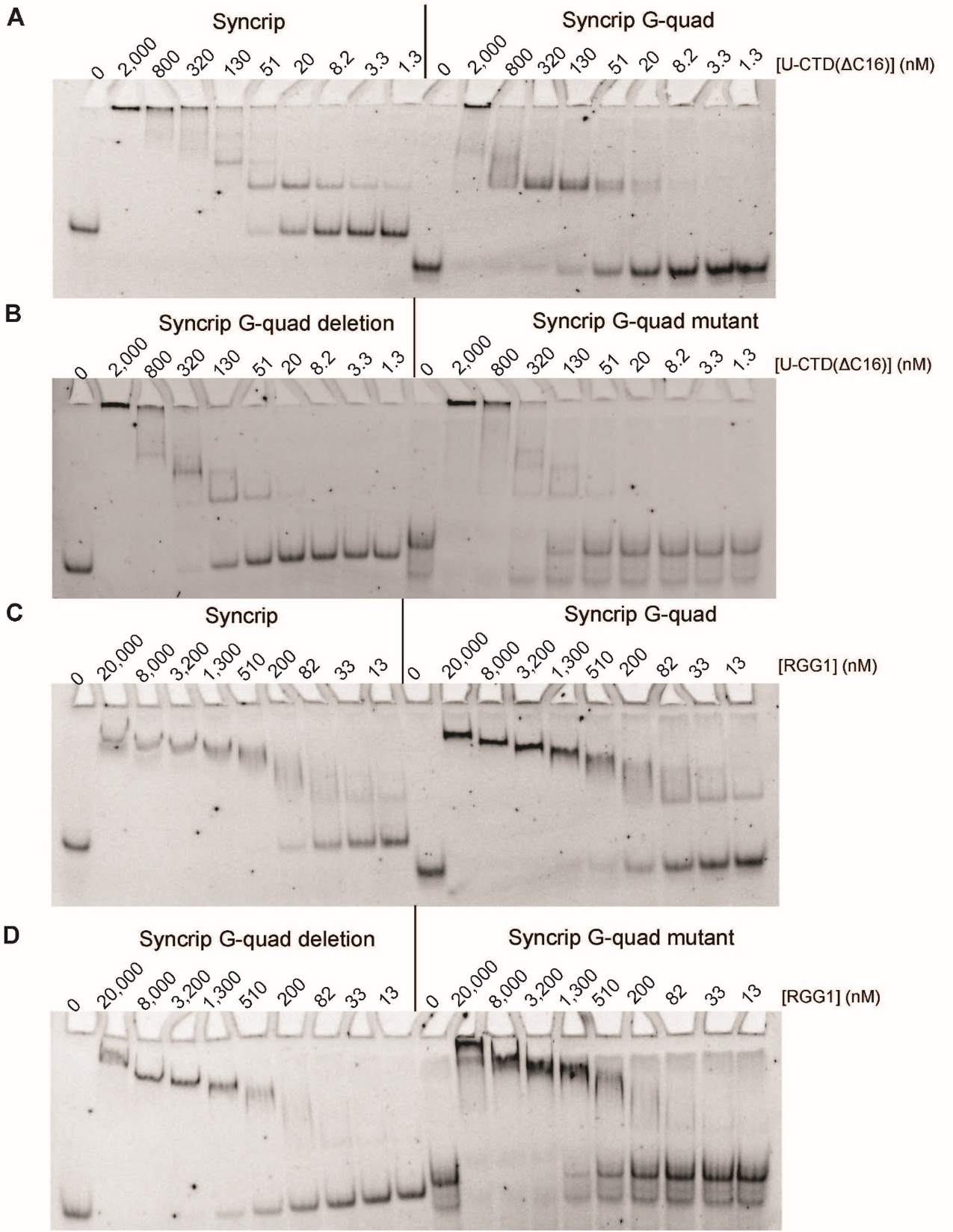

**Figure S3. EMSA analysis of selectivity for RNA G-quadruplexes.** (A) EMSAs depicting U-CTD(∆C16) binding to Syncrip and Syncrip G-quad RNAs. (B) EMSAs depicting U-CTD(∆C16) binding to Syncrip ∆G-quad (left) and Syncrip GtoC (right) RNAs. (C) EMSAs depicting RGG1 binding to Syncrip and Syncrip G-quad RNAs. (D) EMSAs depicting RGG1 binding to Syncrip ∆G-quad and Syncrip GtoC RNAs.

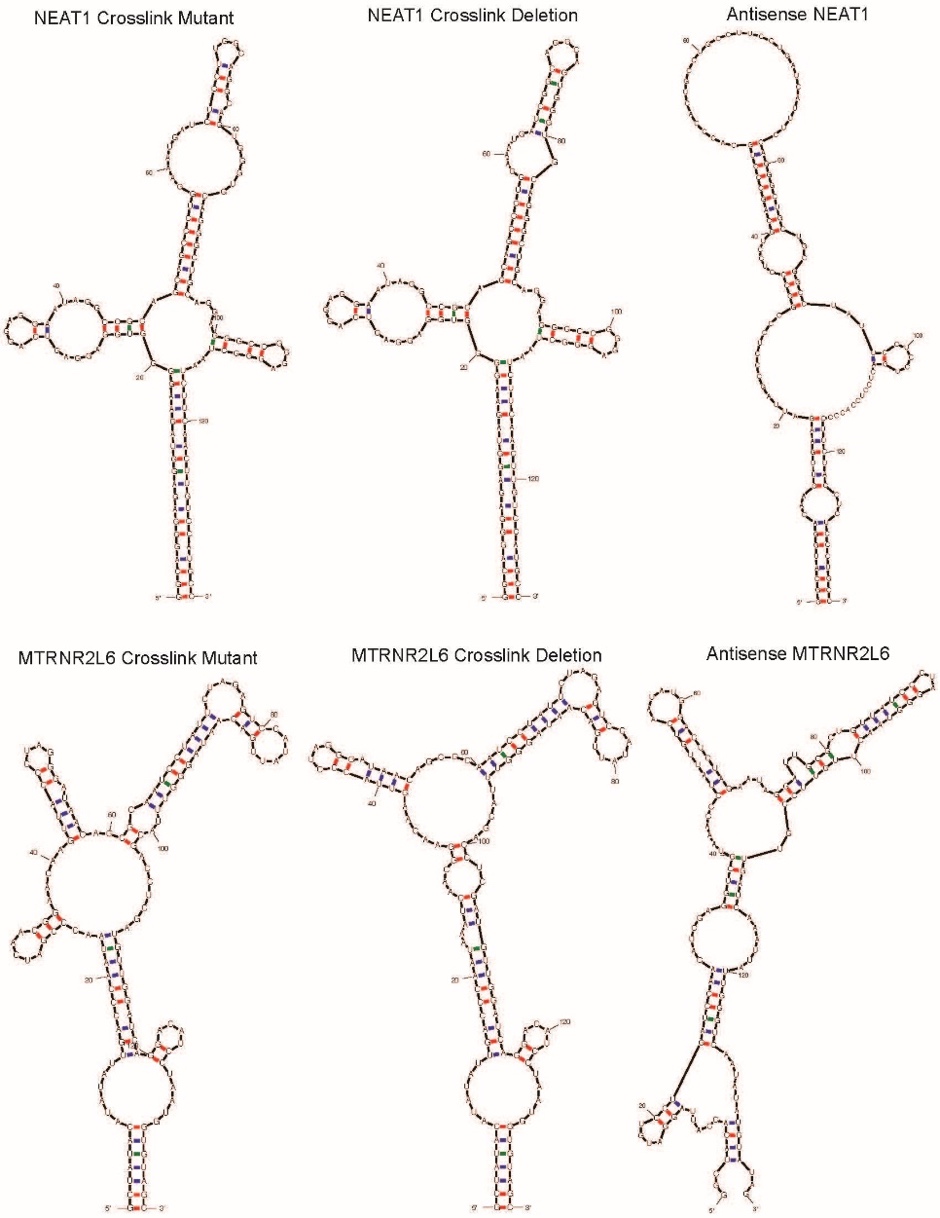

**Figure S4: Mfold structure predictions of NEAT-1 and MTRN2L6 variants.**

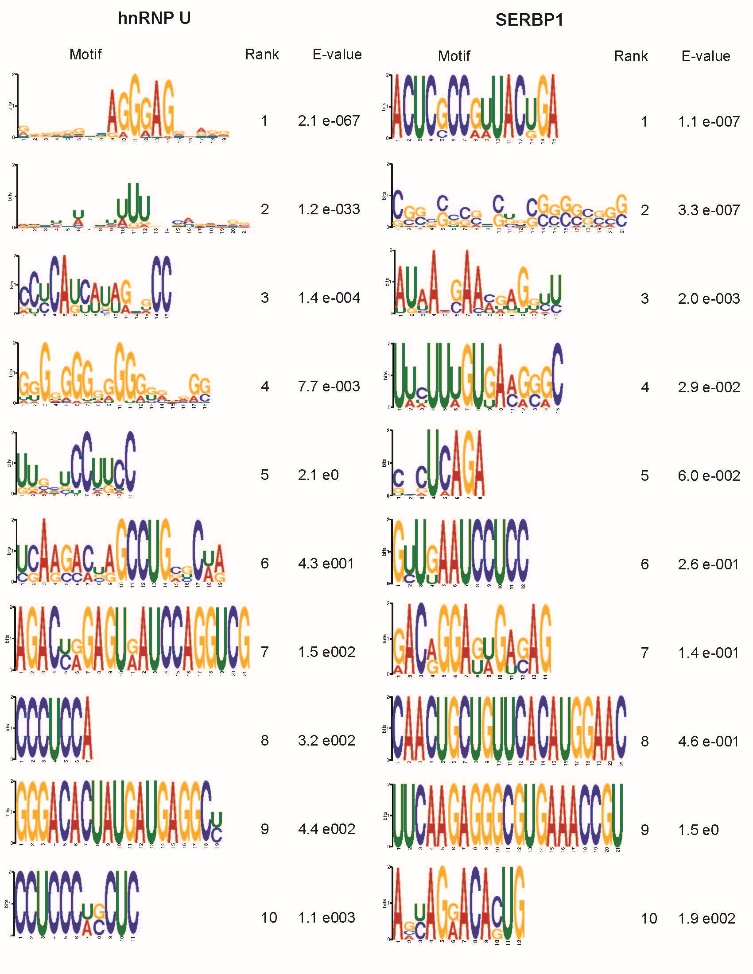

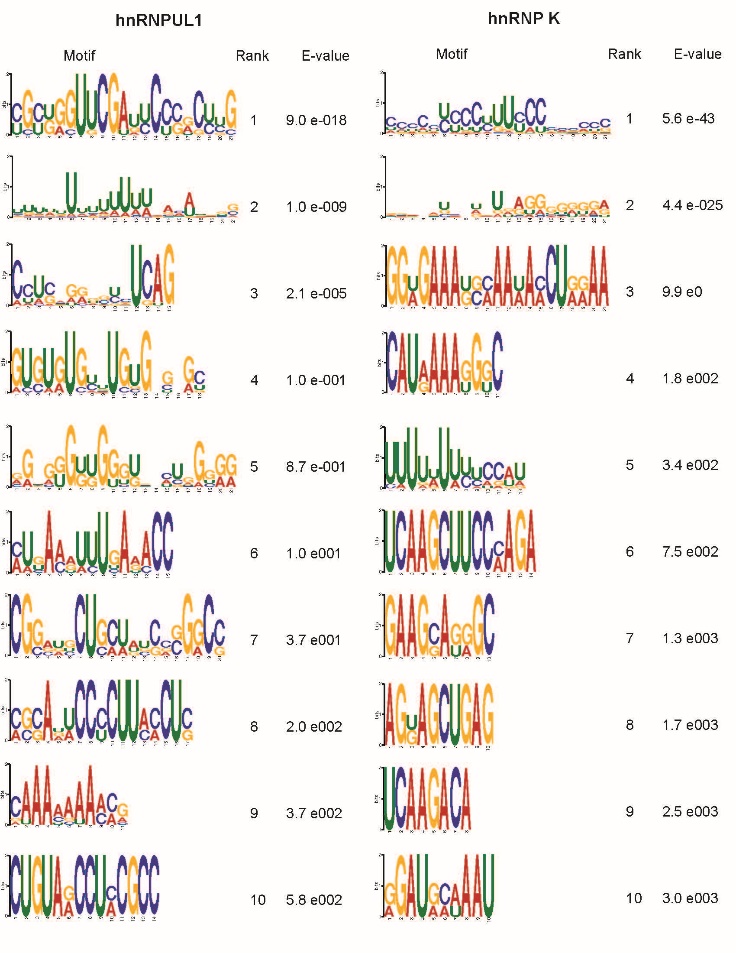

**Figure S5: Top 10 MEME Motifs identified from PureCLIP sites.** Continued on next page.

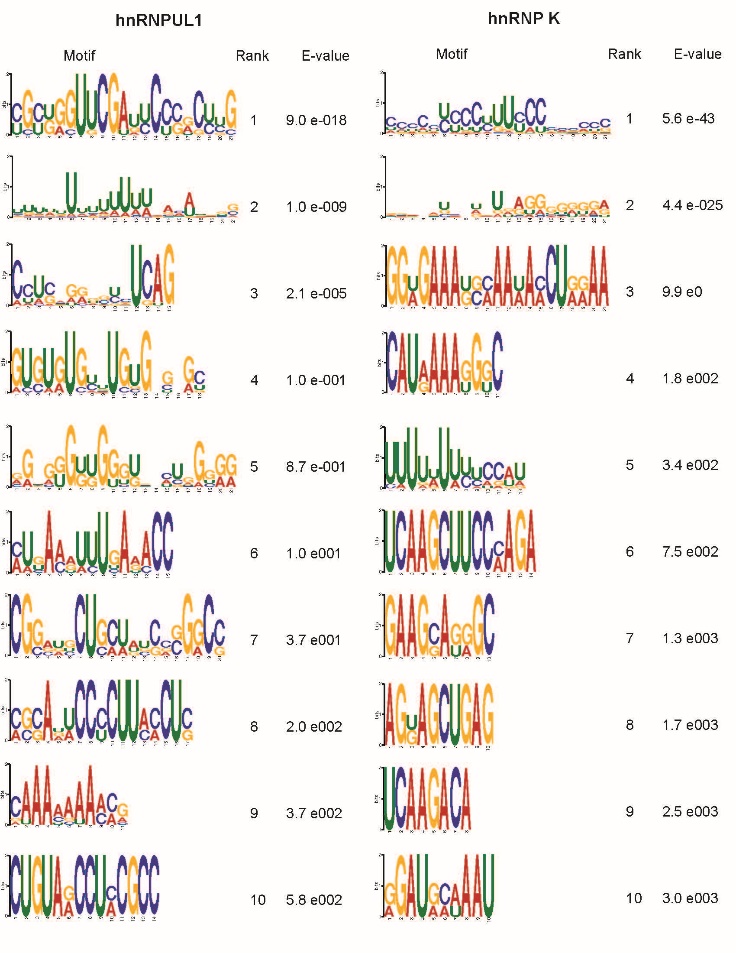

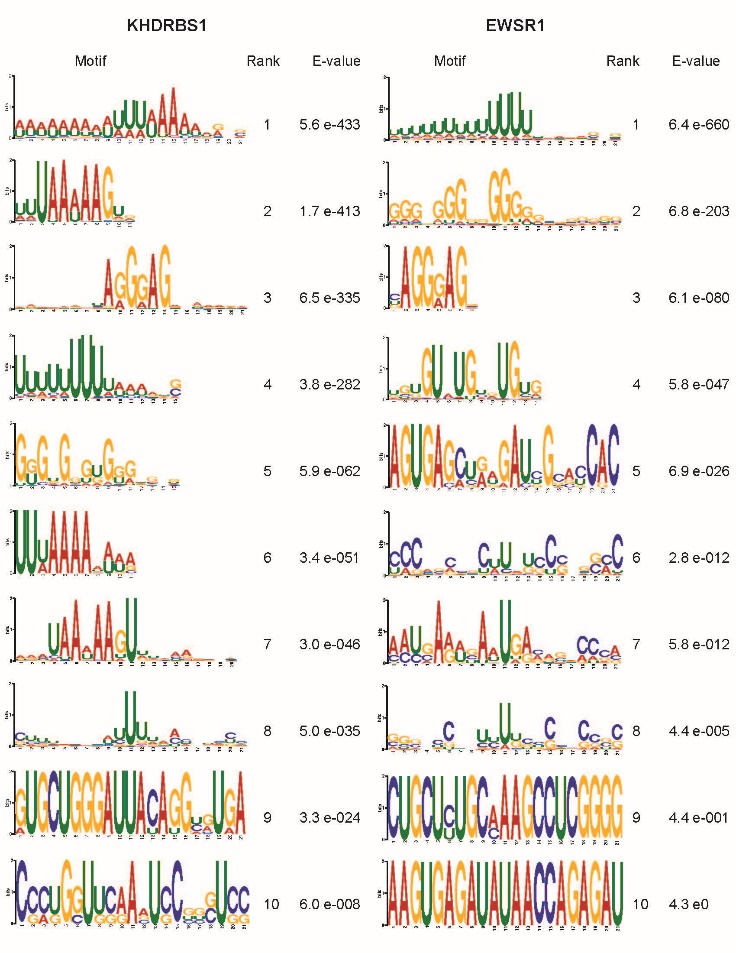

**Figure S5: Top 10 MEME Motifs identified from PureCLIP sites.** Continued on next page.

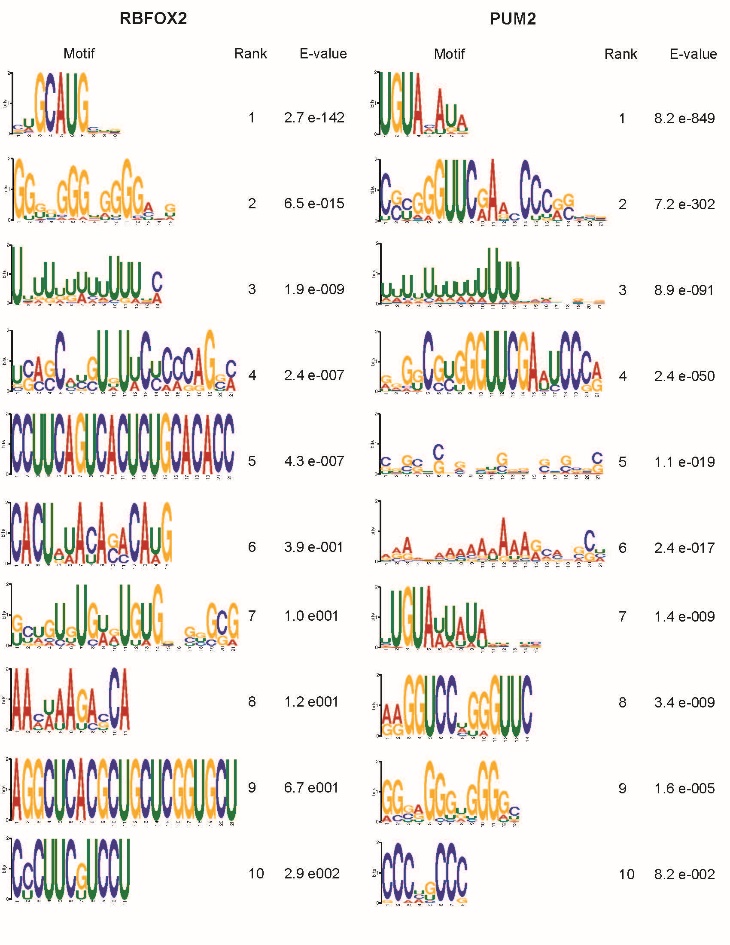

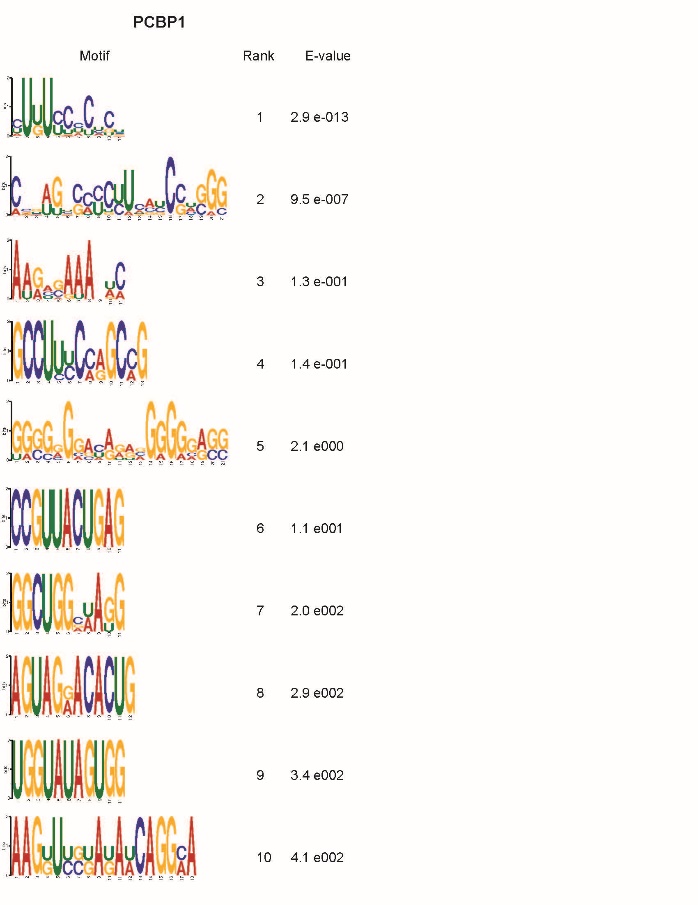

**Figure S5: Top 10 MEME Motifs identified from PureCLIP sites for a variety of RNA-binding proteins.** All eCLIP datasets for the indicated proteins were accessed from the ENCODE Consortium. PureCLIP sites conserved across all available biological replicates were extended 10 nt in each direction and used as input for MEME. The top 10 motifs identified for each protein are listed with their rank and E-value.

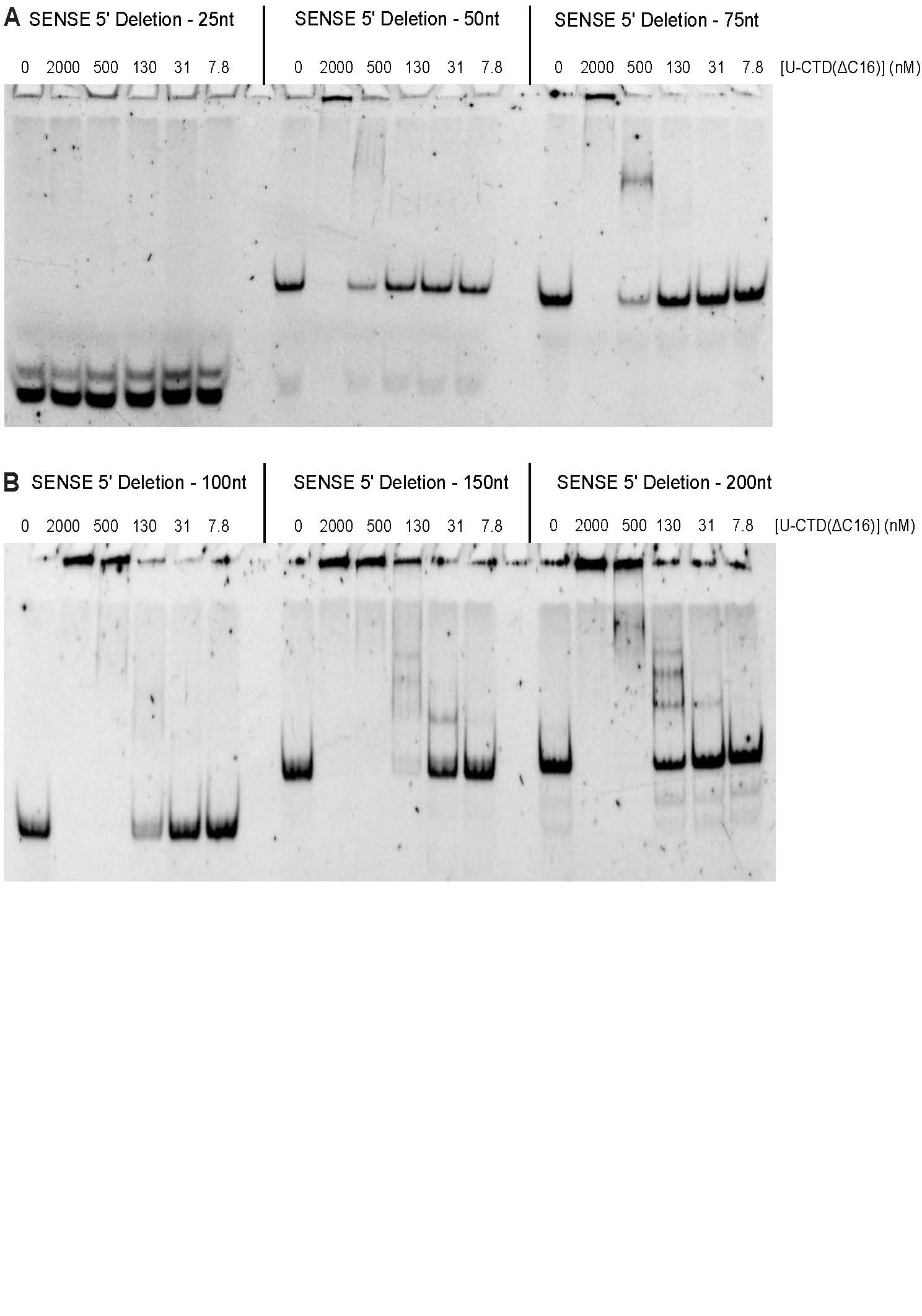
**Figure S6: EMSAs confirm length-dependent binding by hnRNP U.** EMSAs were performed with the U-CTD(∆16) protein fragment. (A) EMSA of the 25-, 50-, and 75-nt RNAs. (B) EMSA of the 100-, 150-, and 200-nt RNAs.

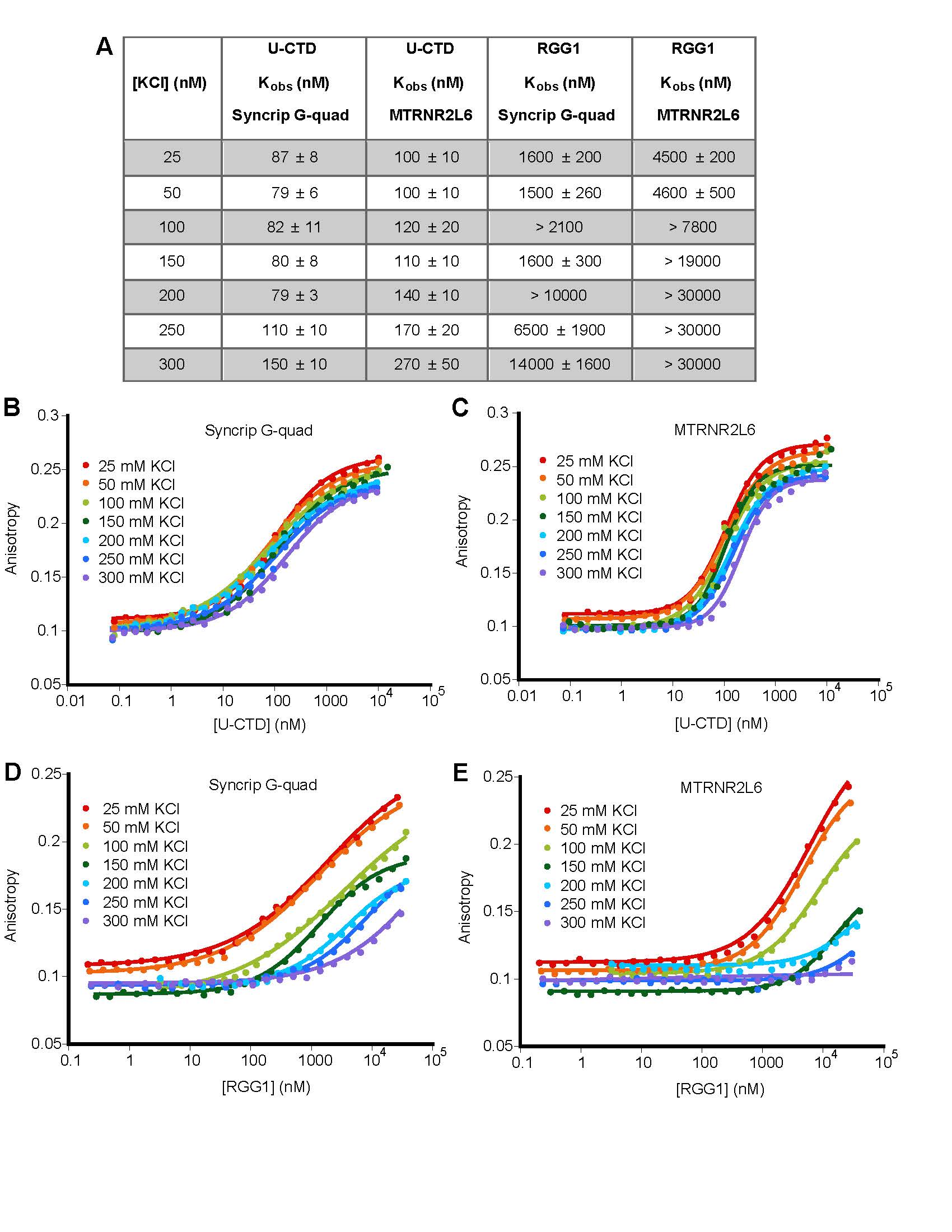

**Figure S7: U-CTD and RGG1 display varying salt dependencies.** (A) Table of FA-derived binding measurements performed at varying KCl concentrations. For each condition, the average K_obs_ and associated SEM are shown. ">" denotes estimated K_obs_ values from unsaturated binding curves. (B) Representative FA curves of U-CTD bound to Syncrip G-quad RNA at increasing [KCl]. (C) Representative FA curves of U-CTD bound to MTRNR2L6 RNA at increasing [KCl]. (D) Representative FA curves of RGG1 bound to Syncrip G-quad RNA at increasing [KCl]. (E) Representative FA curves of RGG1 bound to MTRNR2L6 at increasing [KCl].

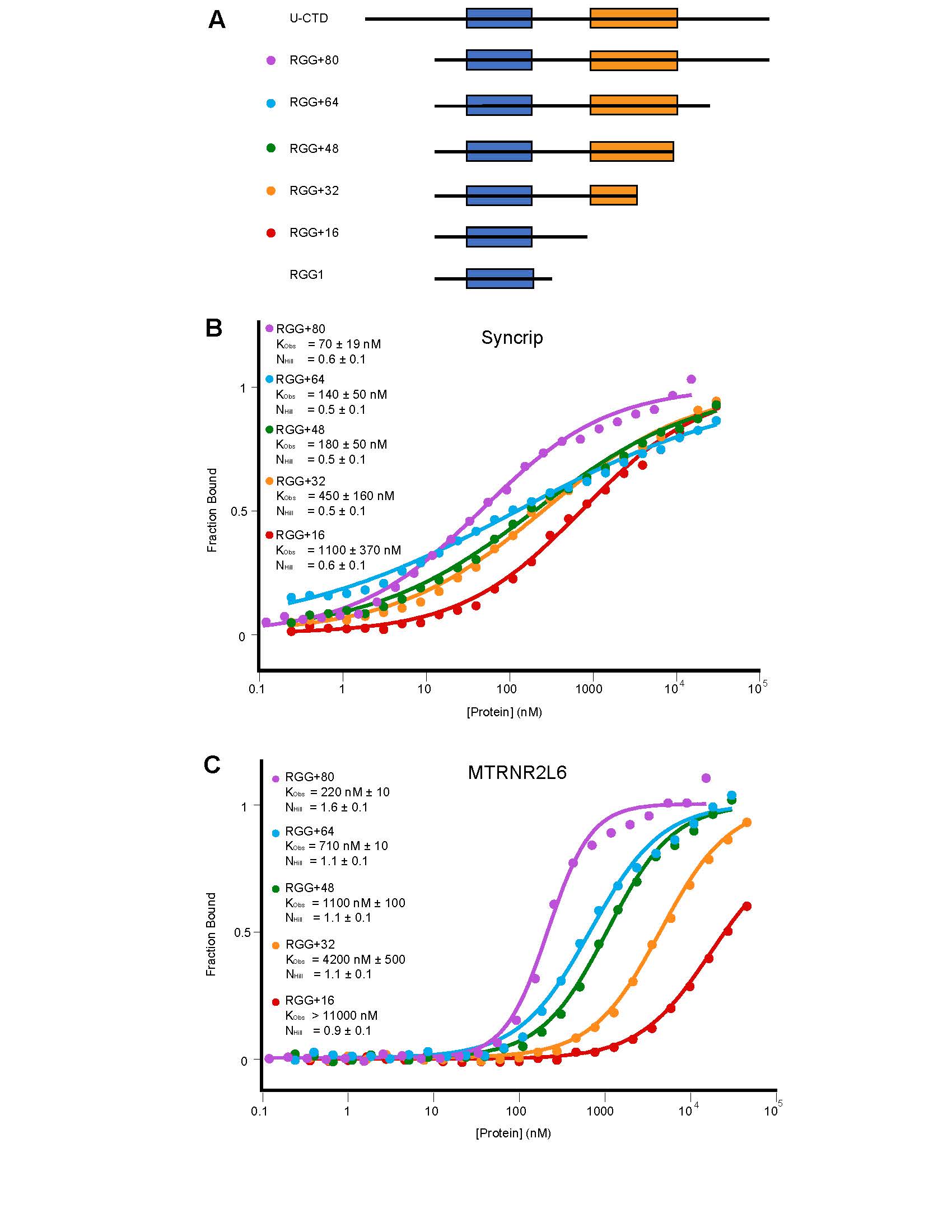
**Figure S8: The full hnRNP U CTD contributes to binding.** (A) Schematic of hnRNP U protein fragments. The blue box represents the canonical RGG/RG motif spanning positions 714-739. (B) Representative normalized FA curves of hnRNP U protein fragments bound to Syncrip RNA, with average Hill coefficients and K_obs_. (C) Representative normalized FA curves of hnRNP U protein fragments bound to MTRNR2L6 RNA, with average Hill coefficients and K_obs_. ">" denotes estimated K_obs_ values from unsaturated binding curves.
